# Fitness Landscape for Antibodies 2: Benchmarking Reveals That Protein AI Models Cannot Yet Consistently Predict Developability Properties

**DOI:** 10.64898/2025.12.27.696706

**Authors:** Michael Chungyoun, Jeffrey Gray

## Abstract

A prominent application of machine learning in therapeutic antibody design is the development of models that can generate or screen antibody candidates with a high probability of success in manufacturing and clinical trials. These models must accurately represent sequence-structure-function relationships, also known as the fitness landscape. Previous protein function benchmarks examine fitness landscapes across diverse protein families, but they exclude antibody data. Here, we introduce the second iteration of the Fitness Landscape for Antibodies (FLAb2), the largest public therapeutic antibody design benchmark to date. The datasets collected in FLAb2 contain developability assay data for over 4M antibodies across 32 studies, encompassing seven properties of therapeutic antibodies: thermostability, expression, aggregation, binding affinity, pharmacokinetics, polyreactivity, and immunogenicity. Using the curated data, we evaluate the performance of 30 artificial intelligence (AI) and biophysical models in learning these properties. Protein AI models on average do not produce statistically significant correlations for most (80%) of developability datasets. No models correlate with all properties or across multiple datasets of similar properties. Zero-shot predictions from pretrained models are incapable of accurately predicting all developability properties, although several models (IgLM, ProGen2, Chai-1, ESM2, ISM, IgFold) produce statistically significant correlations for multiple datasets for thermostability, expression, binding, or immunogenicity. Fine-tuning with at least 10^2^ points improves performance on thermostability, aggregation, and binding, but polyreactivity and pharmacokinetics lack enough data for significance. Yet it is humbling to observe that given enough developability data (10^3^ points), a fine-tuned one-hot encoding model can match the performance of fine-tuned billion-parameter pretrained models. Training data composition influences performance more than model architecture, and intrinsic biophysical properties (thermostability) are more readily learned than extrinsic properties (immunogenicity, pharmacokinetics). Controlling for germline distance with partial correlation reveals that protein language models draw substantially on evolutionary signal; on average, germline edit distance accounts for 40% of their apparent predictive power. FLAb2 data are accessible at https://github.com/Graylab/FLAb, together with scripts that allow researchers to benchmark, compare, and iteratively improve new AI-based developability prediction models.

## 2 Introduction

The innate and adaptive immune system is pivotal for safeguarding the human body, with antibodies employed as specialized proteins evolved to combat diseases. Antibody engineering exploits their therapeutic potential, resulting in over 200 marketed antibody therapeutics and a pipeline of nearly 1,400 investigational product candidates currently undergoing evaluation in clinical studies as treatments for a wide variety of diseases^1^. The efficacy of therapeutic antibody candidates hinges on achieving a delicate balance of drug-like biophysical properties, also known as developability properties, characterized by intricate tradeoffs where enhancing one property may compromise others.

The rapidly advancing field of artificial intelligence is increasingly being applied to antibody design, enabling the generation of novel and diverse therapeutic candidates with favorable developability profiles in significantly reduced timeframes. Generative models can create antibodies in various formats - including VHHs, scFvs, and Fvs - that target a broad range of antigen epitopes, such as those from infectious agents like intestinal pathogens, influenza, RSV, SARS-CoV-2; or endogenous molecular targets like IL-7, TNFα and GPCRs^2–5^. In parallel, pretrained protein language models have been used in an unsupervised, zero-shot manner to guide mutational selection, enhancing the stability and antigen-binding affinity of wild-type antibodies^6,7^. Protein language model embeddings have also been leveraged to fine-tune downstream models for improved prediction of key developability properties, including clearance^8^, viscosity^9^, and in vitro assays^10–12^.

As the diversity of deep learning approaches increases^13,14^, a systematic benchmark is needed for evaluating performance. Current antibody design methods are typically evaluated with native sequence or structure recovery^15–19^, which does not provide a complete indication of therapeutic potential. Other antibody design studies rely on sequence- or structure-based predictors of biophysical properties that have not yet been robustly validated against experimental antibody developability data^20,21^.

Here, we provide a therapeutic antibody design benchmark dataset by curating experimental fitness data from over 30 studies spanning antibody thermostability, expression, aggregation, binding affinity, pharmacokinetics, polyreactivity, and immunogenicity. Our dataset collection, the Fitness Landscape for Antibodies (FLAb2), is the largest publicly available database of diverse developability data covering many relevant therapeutic properties. With FLAb2, we contribute (1) diverse datasets of therapeutic antibody assay data, (2) an assessment of the performance of widely adopted, pretrained and fine-tuned deep learning models for proteins, (3) an interpretation for what protein AI models learn and how model characteristics influence performance, and (4) scripts for zero-shot scoring and fine-tuning pretrained models. After introducing the data, we demonstrate its value by benchmarking a collection of models relevant to antibodies for their ability to correlate likelihoods to fitness properties. Our long-term vision is that FLAb2 will help the development of models that can generate or filter new antibody design candidates more efficiently than what is often done experimentally.

## 3 Related work

### 3.1 Several databases provide antibody sequence and structure data, but not developability

The Observed Antibody Space (OAS)^22^ and NaturalAntibody AbNGS^23^ collectively provide 5B annotated sequences from immune repertoires. The Structural Antibody Database (SAbDab)^24^ contains 10K antibody structures from the Protein Data Bank (PDB) and is continually updated. SAbDab provides over 500 experimental binding affinity data points associated with some of the antibody-antigen crystal structures, but does not include other developability properties.

### 3.2 Most protein fitness benchmarks exclude antibodies

Previous endeavors to establish benchmarks for function prediction have laid a foundation for protein engineers to assess new models. The Critical Assessment of Functional Annotation (CAFA)^25^ tests the assignment of gene ontology classes to proteins. The Task Assessing Protein Embeddings (TAPE)^26^ evaluates different pretrained models in predicting three protein structure properties (remote homology, secondary structure, residue contacts), as well as two fitness properties (fluorescence and stability). The Fitness Landscape for Proteins (FLIP)^27^ examines the performance of masked language models, convolutional networks, and evolutionary models at expression, binding, and thermostability prediction. ProteinGYM^28^ tests zero-shot and few-shot benchmarking of protein language models from deep mutational scans. Garcia et al.^29^ curated a dataset of 614 experimentally characterized de novo designed monomers and benchmarked the ability of AlphaFold, ProteinMPNN, and ESMFold to zero-shot predict expression and solubility. RemoteFoldSet benchmarks the structural awareness of structure-informed language models^30^. These benchmarks provide insight in the capabilities of AI protein models for protein function prediction, but they all exclude antibody data. The absence of a publicly accessible benchmark dataset for therapeutic antibodies hinders the ability to identify which model properties contribute to optimal performance in predicting antibody fitness landscapes.

### 3.3 Existing antibody benchmarks focus on binding prediction

Antibody binding datasets have been curated for benchmarking. AstraZeneca used publicly available binding data from AbSci^31^, Porebski et al.^32^, and 85 data points generated in-house to benchmark their antibody sequence ranking model DiffAbXL against other approaches^33^. AbRank^34^ benchmarked five deep learning models at antibody binding affinity prediction, using a database of 300K antibody-antigen pairs curated from nine studies. NaturalAntibody curated AbDesign^35^, a dataset of 14 antibody-antigen complexes, each paired with mutational data totaling over 600 affinities. AbAgym^36^ curated 67 deep mutational scan datasets to benchmark ML models at predicting the change in antibody-antigen binding affinity upon mutation. AbBiBench^37^ curated 180K experimental binding measurements and benchmarked protein AI models at antibody binding affinity maturation. These benchmarks assess binding affinity, a key step in therapeutic design, but an antibody’s success also hinges on other developability factors. Our preliminary version of FLAb^38^ provided 30K datapoints across six developability properties to benchmark six deep learning models at zero-shot fitness prediction. Here, our updated release, FLAb2, integrates the previously described datasets with additional datasets mined from publications, yielding a database of 4M antibodies, and we benchmark 30 protein AI models on both zero-shot and few-shot prediction.

## 4 Fitness landscape data

Jain et al.^39^ defines desirable developability characteristics as (1) high conformational stability, (2) high-level expression, (3) low propensity for aggregation, (4) high binding affinity towards the target antigen, (5) low polyreactivity, and (6) low immunogenicity (**Fig. 1**). FLAb2 includes sequences and measurements from Jain et al. and 31 other studies, which we use to assess the efficacy of protein AI models in capturing essential characteristics of therapeutic antibodies (**Fig. 2**, **Table S1**). We collected *in vitro* and *in vivo* assays commonly used in therapeutic antibody discovery campaigns and summarized their counts by developability category and define each fitness landscape in **Fig. S1** and **Supp. Section 10.2**, respectively. Each sequence is associated with at least one fitness label pertaining to the developability properties. Most of the collected datasets contain measurements from a single assay for a relatively small set of antibody sequences, with only a few datasets providing multiple assays per-sequence and large sequence sets (**Fig. 2**).

**Figure 1:**
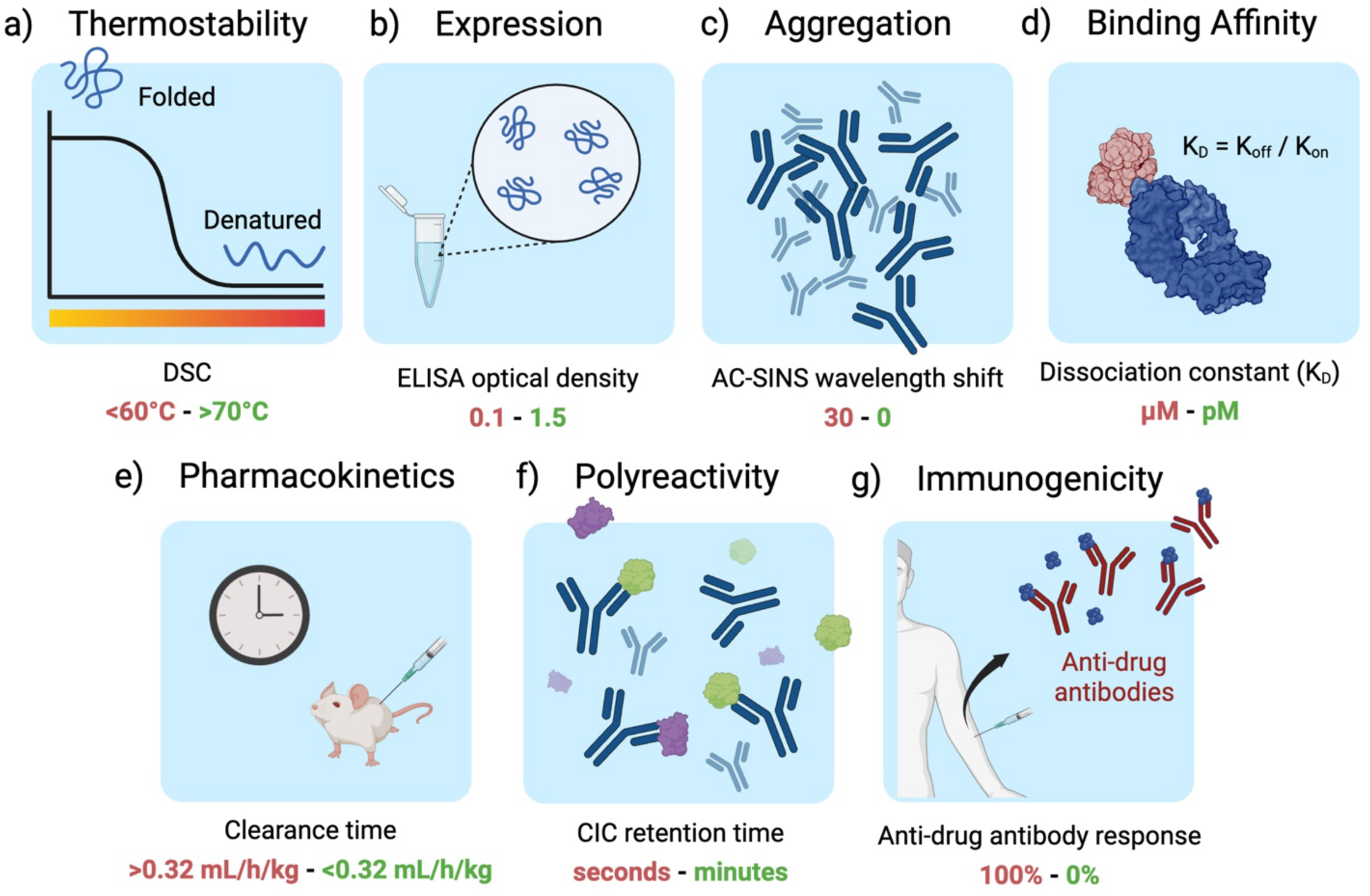
Seven classes of biophysical data are relevant to antibody developability. Left to right: Increasing machine learning prediction complexity. Provided below each category is an example assay used to quantify the property, with desirable values shown in green and undesirable values in red. (a) Antibody thermostability (Tm) is assessed through differential scanning calorimetry (DSC), which measures the heat capacity change of a sample with temperature; higher Tm values are considered better. (b) Expression is often assessed using enzyme-linked immunosorbent assay (ELISA) fluorescence with high optical density indicating desired high expression. (c) Aggregation can be measured using affinity-capture self-interaction nanoparticle spectroscopy (AC-SINS), where no change in the measured plasmon wavelength shift is ideal. (d) Binding affinity to a target antigen is assessed using the equilibrium dissociation constant (K_d_), with a desirable K_d_ typically falling in the low nanomolar to picomolar range; smaller K_d_ values indicate a desired tighter binding. (e) Pharmacokinetics is measured by timing the clearance of a drug *in vivo* through an animal, typically mice or cynomolgus monkeys, where a low clearance rate is preferred. (f) Polyreactivity can be measured with cross-interaction chromatography (CIC) retention time, where therapeutic antibodies are expected to have a low retention time in the column indicating minimal non-specific interactions. (g) Immunogenicity is quantified as the percentage of patients developing anti-drug antibodies (ADAs) after therapeutic administration, where an ideal, non-immunogenic antibody results in 0% ADA response within a patient population.

**Figure 2:**
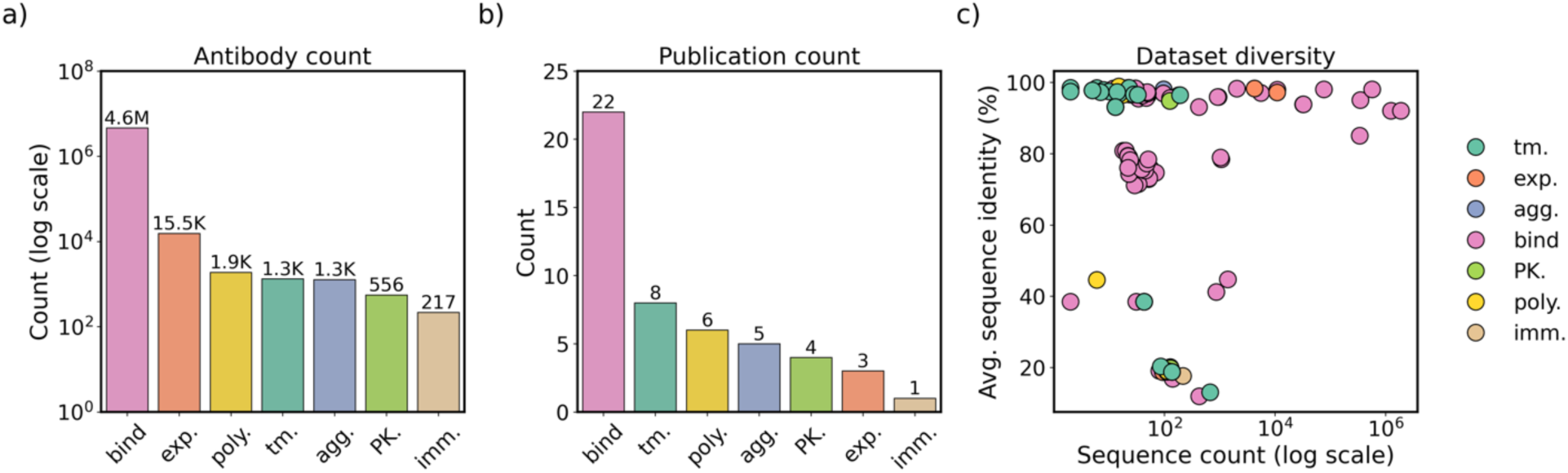
FLAb2 is composed of developability data collected from publications, competitions, and collaborators. (a) Most of the collected data is for binding, and data for pharmacokinetics is scarce. (b) Binding data is gathered from 22 publications. Except for binding, in each category, we were only able to identify 8 or fewer studies with open data. (c) Most of the datasets are either large (>1,000 sequences) but closely related (e.g. from a deep mutational scan of single point mutations) or small (<200 sequences) and diverse (antibodies from unique clonotypes).

## 5 Pipeline for model evaluation

Protein models that produce statistically significant correlations with the antibody fitness landscapes can be reliable predictors for new therapeutic antibody design candidates. To benchmark protein models, we created a pipeline to run prediction models and correlate them to each experimental dataset (**Fig. 3**).

**Figure 3:**
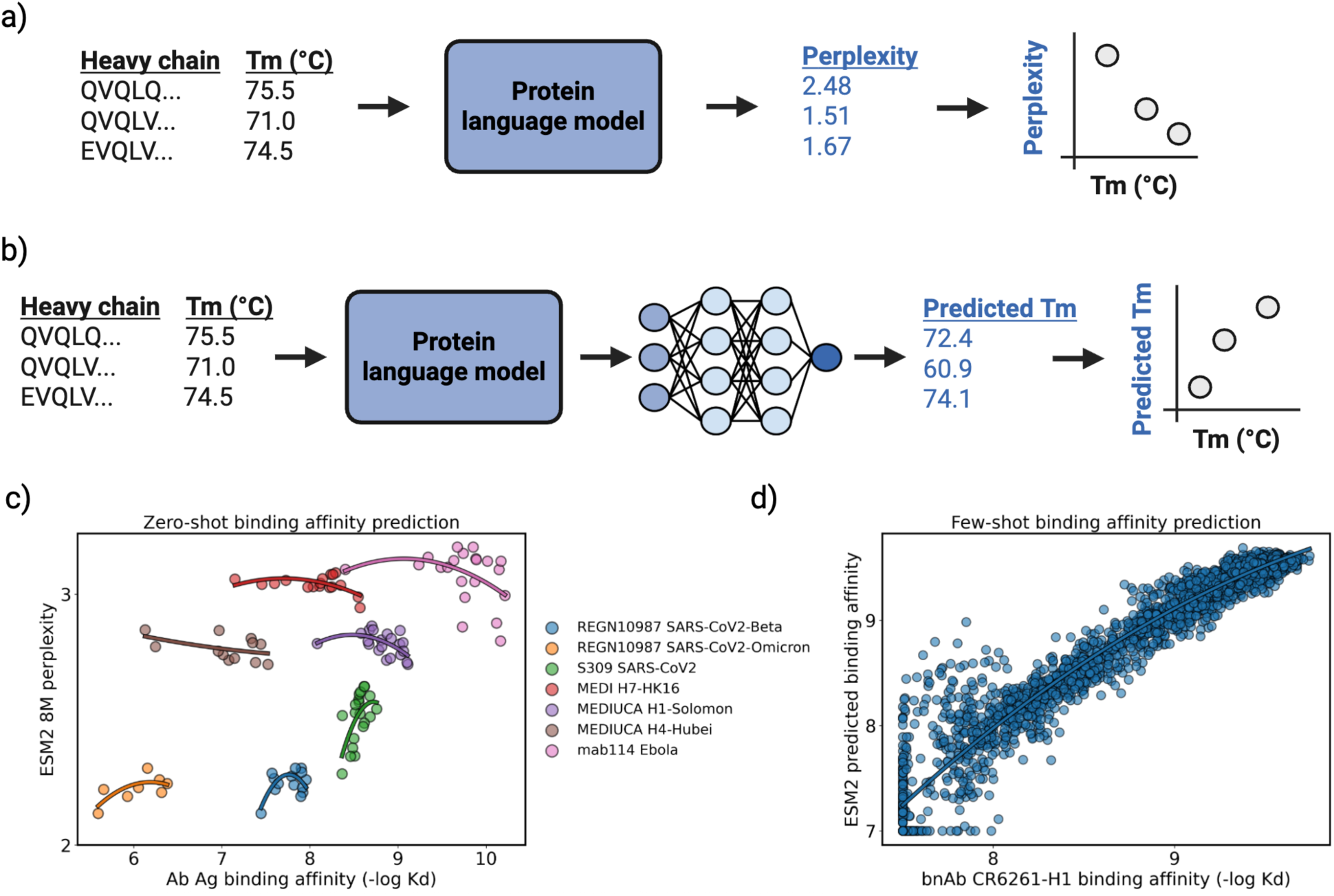
Pipeline for benchmarking protein AI models. All fitness datasets contain columns for antibody heavy chain sequence, antibody light chain sequence, and an associated fitness metric. (a) Zero-shot benchmarking: For each protein language model, we separately input the heavy and light sequence to return two perplexity scores, and we tabulate the average perplexity between the two sequences. For structure-conditioned language models, we first predict the antibody structure with Chai-1^39^, and then tabulate the single perplexity scored from the model. Spearman’s correlations are calculated between average perplexity and the fitness measure. (b) Few-shot benchmarking: For each protein language model, we separately input the heavy and light chain sequences to return two embedded representations. The pairs of embeddings are concatenated, pooled, and used to train a two-layer fully connected network for regression with an 80/10/10 train/validation/test split. Spearman’s correlations are calculated between predicted fitness and true fitness. No antigen information is provided for any benchmarked models. We use Spearman’s correlation between model outputs and true fitness labels to capture any monotonic relationship between variables and to account for the non-normally distributed data we use to benchmark. Some assay labels were inverted to ensure that, in all cases, a positive Spearman’s correlation indicates better alignment between the model’s confidence and the assay values. We provide examples for (c) zero-shot and (d) few-shot benchmarking for antibody binding prediction.

### 5.1 Zero-shot benchmarking

Zero-shot antibody fitness prediction is the task of using a model pretrained on protein sequence and structure to score a new set of antibodies, where the score is assumed to be descriptive of the antibody’s fitness, even though the model is not explicitly trained on this task. For zero-shot benchmarking (**Fig. 3a**), we use the antibody variable region sequence or structure as inputs for each model and output perplexity scores (averaged over all residues in the heavy and light chains). We use perplexity because it represents the confidence that the new antibody is similar to the learned distribution of high-fitness proteins from the pretraining dataset. If a protein language model implicitly captures the biophysical landscapes of antibodies, it should assign high confidence (low perplexity) to previously unseen high-fitness antibodies and low confidence (high perplexity) to low-fitness antibodies (examples in **Fig. 3c**). In zero-shot testing, all models were previously trained in their respective studies - we performed no additional fine-tuning prior to calculating perplexities.

We report the Spearman’s correlations and associated p-value to establish the connection between the model uncertainty values and the fitness metrics associated with the sequences in the dataset. An optimal zero-shot predictor will display a Spearman’s correlation close to 1.0 between model perplexity and developability labels. In contrast, a Spearman’s correlation of 0.0 suggests no relationship between these metrics, while a correlation of -1.0 indicates an inverse relationship between the model’s predictions and the developability label.

**Fig. 3c** provides an example for zero-shot benchmarking. Ideally, the model should be more confident in the higher binding affinity antibody than the variants with lower binding affinity. This correct downward trend is observed for several candidates (MEDIUCA H1-Solomon, ρ = -0.53; MEDIUCA H4-Hubei, ρ = -0.47; mab114 Ebola, ρ = -0.28; MEDI H7-HK16, ρ = -0.20), yet the model incorrectly displays more confidence with the weaker affinity antibodies for S309 SARS-CoV2 (ρ = 0.57) and REGN10987 SARS-CoV2-Omicron (ρ = 0.57).

### 5.2 Few-shot benchmarking

Few-shot antibody fitness prediction is the task of supervised fine-tuning a protein AI model on a developability dataset for predicting that property. For few-shot benchmarking (**Fig. 3b**), we explore an embedding-based fine-tuning approach. We use each protein language model to generate an embedded representation for each antibody and train a regression model on part of the developability dataset to predict developability of the holdout set. Predictions on the test set are evaluated based on the Spearman’s rank order correlation between the predicted and true assay values. Like the zero-shot setting, few-shot benchmarking is performed per dataset, and statistical significance is evaluated based on the associated p-value of the Spearman correlation. If a protein language model captures the biophysical landscapes of a pretraining set of antibodies or general proteins, the rich features in extracted embeddings should allow a learnable mapping between embeddings and developability properties. Like zero-shot benchmarking, an ideal predictor will have a Spearman’s correlation approaching 1.0 between predicted developability and true developability.

In the few-shot benchmarking example in **Fig. 3d**, there is a significant (p-value = 0.00) positive correlation (ρ = 0.93) between the predicted and true fitness label for the strong binders of the broadly neutralizing antibody CR6261 in the upper right-hand quadrant, while showing relatively more noise for predicting weak binders in the lower left-hand quadrant.

### 5.3 Models benchmarked

We benchmark a diverse set of AI models associated with antibody design: (1) Decoder-only language models are trained using next-token prediction, and we investigate IgLM^40^ and ProGen2^41^; (2) encoder-only language models capture continuous representations of sequences, and we investigate AntiBERTy^42^, ESM-2^43^, and ISM^44^; (3) inverse folding models predict protein sequences from structure, and we investigate AbMPNN^45^, ESM-IF^46^, and ProteinMPNN^47^; (4) structure prediction models predict protein structures from sequence, and we investigate IgFold^48^ and Chai-1^49^. As a baseline for physics-based approaches, we calculate PyRosetta^50^ total energy and BioPython^51^ charge terms. We provide a more in-depth description of each model in **Table S2** and further details about the categories of benchmarked models in **Supp. Section 10.3**.

## 6 Zero-shot results

We evaluated whether model likelihoods, expressed as average perplexities, correlate with experimentally measured antibody developability properties.

### 6.1 Most models do not have statistically significant correlations with most datasets

We first investigate how many statistically significant (p-value < 0.05) Spearman’s correlations each zero-shot model has to each developability dataset. The structure-informed masked language model ISM 650M displayed the highest count of statistically significant correlations, with 94 correlations or 39% of the total developability data sets (**Fig. 4a**). In order of decreasing count, protein AI models on average displayed significant correlations to 51% of immunogenicity datasets (**Fig. 4h**), 35% of expression datasets (**Fig. 4c**), 24% of thermostability datasets (**Fig. 4b**), 20% of binding datasets (**Fig. 4e**), 17% of aggregation datasets (**Fig. 4d**), 16% of polyreactivity datasets (**Fig. 4g**), and 14% of pharmacokinetics datasets (**Fig. 4f**). The observed variation in the number of statistically significant correlations across developability categories likely reflects a combination of factors that are difficult to disentangle, including whether a model captures the underlying biophysical property, whether a data set is simply too small, and whether the biological assay readouts are too noisy. Some datasets exhibit statistically significant correlations across multiple models, suggesting that some data sets are clean and large enough to be used to fairly benchmark models. Accordingly, we designated a subset of datasets that show at least five statistically significant model–dataset correlations for further comparison (**Table S3**); the corresponding distributions of Spearman’s correlation coefficients for each model are shown in **Fig. 5**.

**Figure 4:**
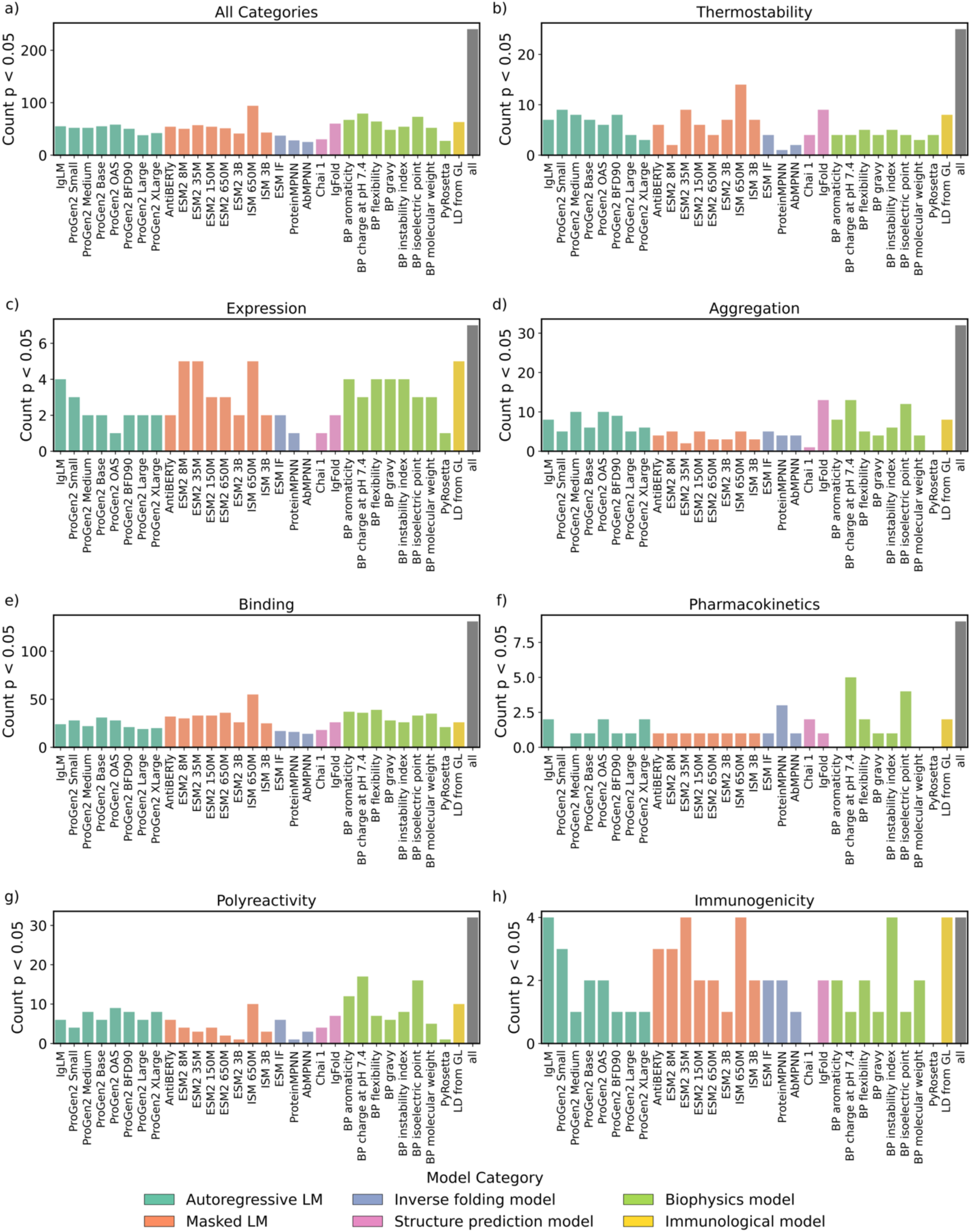
Count of significant correlations per zero-shot model. Count of significant (p-value < 0.05) correlations each protein AI and physics-based model had within each developability category. The “all” column within each subplot represents the total number of datasets within that category.

**Figure 5:**
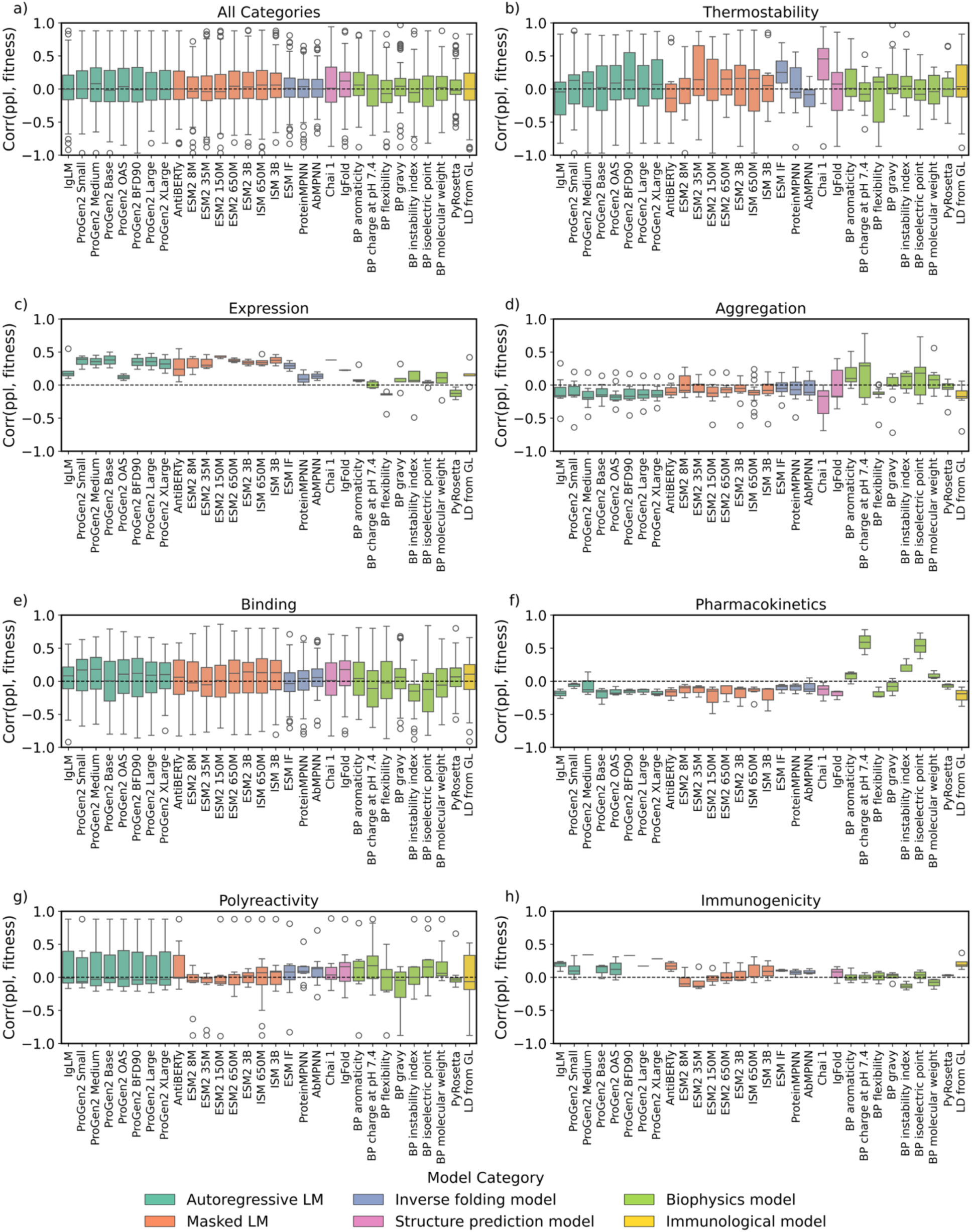
Distribution of zero-shot prediction performances for each model. Models are colored based on their architecture, and the *y*-axis displays the range of Spearman’s correlations between model confidence and developability label, where a correlation approaching 1.0 is ideal. Some of the assay labels are inverted so that a positive correlation between fitness and assay value always indicates improved model performance. We only report Spearman’s correlations with datasets that exhibit at least five significant dataset-model p-values to ensure our conclusions are statistically sound.

### 6.2 No single model predicts all developability properties with high accuracy

Within each developability category, we investigate the average performance across all protein AI models and identify the best performing model. Expression was the least challenging task (average Spearman’s rank correlation ρ = 0.22), whereas aggregation was the most difficult (ρ = -0.07) (**Table S4**). From easiest to hardest, the average task ranking across all models was: expression > immunogenicity > thermostability > binding > polyreactivity > pharmacokinetics > aggregation.

IgFold was the best overall performer (ρ = 0.12) (**Fig. 5a**), and most developability categories had a unique top-performing model (**Fig. 5b-g**, **Table S5**). The structure prediction model Chai-1 best predicted thermostability (ρ = 0.45) (**Fig. 5b**); the masked language model ESM2 150M best predicted expression (ρ = 0.44) (**Fig. 5c**); BioPython charge at pH 7.4 best predicted aggregation (ρ = 0.29) (**Fig. 5d**), pharmacokinetics (ρ = 0.59) (**Fig. 5f**), and polyreactivity (ρ = 0.18) (**Fig. 5g**); and the autoregressive language model ProGen2 Medium best predicted immunogenicity (ρ = 0.34) (**Fig. 5h**, **Table S5**). Top performance in one category did not generalize to others. For instance, while Chai-1 excelled at thermostability prediction, it performed poorly in aggregation prediction (ρ = -0.17). Similarly, ESM2 150M, despite being the best model for expression, ranked as one of the lowest for pharmacokinetics (ρ = -0.15). An ideal antibody developability oracle would exhibit a ρ = 1.0 correlation between model confidence and experimental label for all developability categories, accurately predicting properties across all seven fitness domains. However, no single model achieves uniformly high performance.

Developability properties vary in their predictability using the antibody alone, because some are intrinsic to the antibody sequence and structure, while others depend on extrinsic biological or environmental factors that are not provided to the models. Intrinsic properties such as thermostability and aggregation can, in principle, be inferred from the sequence or fold of the antibody variable domain, whereas extrinsic properties—such as polyreactivity, binding, immunogenicity, expression, and pharmacokinetics—depend on contextual information beyond the sequence. Binding depends on the antigen sequence and structure; immunogenicity depends on the patient’s immune system response; expression depends on host cell machinery; and pharmacokinetics depends on how the body traffics and metabolizes the administered drug. The unexpected difficulty of thermostability prediction may be explained by complications in identifying which portion of the antibody format melts first and is reported in melting curves^52^. These results for antibodies are similar to the results of the general protein benchmark ProteinGYM^28^ in that there is a unique top performing model depending on assay function type, yet different in that some FLAb2 categories had few good predictors (i.e. few models presented a ρ < 0.3), whereas all assay function types in ProteinGYM had multiple models with ρ > 0.4.

### 6.3 Architecture does not significantly impact zero-shot performance

We next ask whether model architecture influences zero-shot performance. Across all developability categories, autoregressive language models (ρ = 0.01), masked language models (ρ = 0.01), inverse folding models (ρ = 0.02), and structure predictors (ρ = 0.07) performed similarly (**Table S6**). Variation in performance is seen between individual developability categories. Autoregressive models performed best at binding and immunogenicity; masked language models performed best at expression; inverse folding models performed best at polyreactivity (tied with structure predictors); and structure predictors performed best at thermostability and polyreactivity (**Table S6**). When comparing two models trained on the same 550M antibody sequence dataset from OAS, AntiBERTy (a masked language model) and IgLM (an autoregressive language model), we found both performed equally in most categories (**Table S5**). Similarly, when comparing the ESM2 suite of masked language models with the ProGen2 suite of autoregressive language models, both performed equally in most categories (**Table S5**). Thus, regardless of whether next-token or masked prediction schemes are used to learn a fitness landscape, models of similar size with identical input representation and training set learn the same data distribution.

### 6.4 Incorporating structure improves zero-shot performance over sequence-only models

We next asked whether learned protein representation influences zero-shot performance. Structure-informed masked language models, inverse folding models, and structure predictors–all of which learn over the protein sequence and structure space–dominated the developability prediction categories (**Table S7**). Comparing the ESM2 suite with the ISM suite allows us to compare two masked models with the same architecture, differing only in the tokenizer: ESM2 tokenizes sequence, while ISM also distills structural representations of the input protein. For thermostability prediction, ISM 3B (ρ = 0.38) outperformed ESM2 3B (ρ = 0.34), however ESM2 650M (ρ = 0.37) outperformed ISM 3B (ρ = 0.34) (**Table S5**). On average, across all categories, the ISM models outperformed the ESM models (**Table S5**). Structural reasoning allows these models to explicitly capture geometric and energetic constraints that sequence-only models overlook. Our findings align with those of Notin et al.^28^, who reported that incorporating structure substantially improves zero-shot prediction relative to sequence-only representations.

### 6.5 Antibody-specific and general protein models excel in different developability domains

We next assessed how models trained specifically on antibody sequences compare to general protein models. General protein models were better at predicting expression (ρ = 0.32), while antibody specific models were better at predicting immunogenicity (ρ = 0.11) (**Table S8**). A direct comparison reveals a slightly different story: Between ProGen2 Medium (trained on UniRef) and ProGen2 OAS (trained exclusively on antibody sequences), both of which are the same autoregressive architecture with 764M parameters, ProGen2 Medium was the better performer for expression (ρ = 0.36), binding (ρ = 0.18), and immunogenicity (ρ = 0.34) (**Table S5**). The different trend when comparing ProGen2 Medium and ProGen2 OAS may be particular to the ProGen2 suite–the original ProGen2 paper also noted that the general protein ProGen2 models were better than ProGen2 OAS for antibody binding, thermostability, and expression prediction^41^. Antibody-specific and general protein models appear to specialize in different aspects of developability. Antibody-specific models perform better at predicting binding and immunogenicity, reflecting their learned understanding of antibody-specific sequence patterns and human-like framework regions. In contrast, general protein models, trained on broader sequence and structural diversity, perform better at predicting expression and thermostability—properties governed by general protein folding and biophysical behavior.

### 6.5 Physics-based and AI-based models capture orthogonal properties of proteins

We then examined the performance of physics-based models for developability prediction and compared it to that of zero-shot protein AI models. PyRosetta was not a top performer in any developability category and generally had no strong correlations to the developability categories (**Table S5**). Several sequence-based BioPython models performed exceptionally well: Across all physics-based and AI-based approaches, BioPython charge at pH 7.4 performed best at aggregation (ρ = 0.29), pharmacokinetics (ρ = 0.59), and polyreactivity (ρ = 0.18) prediction (**Table S5**). Physics-based models exhibited statistically significant correlations across the largest number of models for aggregation, pharmacokinetics, and polyreactivity prediction (**Fig. 5d, f, g**). While several physics-based models performed well at aggregation, pharmacokinetics, and polyreactivity prediction, protein AI models generally performed better at binding, expression, thermostability, and immunogenicity prediction. A charge-based model performs well at aggregation and pharmacokinetic prediction, consistent with the fact that these properties are driven by charged surface patches ^53,54^.

### 6.6 Larger protein AI models do not outperform medium-sized models

We next asked whether the size of the model, determined by the number of parameters, influences zero-shot prediction performance. **Fig. S2** shows that increasing model size did not systematically improve zero-shot performance across categories. For aggregation, binding, and pharmacokinetics, performance remained flat. Thermostability and immunogenicity showed marginal improvement with model scale, while polyreactivity displayed high variability. A common claim with large language models is that scale improves zero-shot performance^55–57^, yet we do not observe this with therapeutic antibody engineering tasks. Notin et al.^28^ similarly found that billion parameter protein language models underperformed million parameter models. This trend may reflect the balance between model capacity and data coverage: overly large models risk memorization, while smaller models generalize more efficiently. The ratio of training tokens to model parameters, as noted by Hou et al.^58^, may provide a more meaningful predictor of generalizability.

### 6.7 Protein AI models are biased towards antibody germline signal rather than developability

Intrigued by the observation of Olsen et al.^59^ that antibody language models have a germline bias, we investigated to what extent protein AI models are correlated to germline signals, rather than the actual developability label. We calculated the Spearman’s correlation between model confidence and the Levenstein distance of the antibody from its respective germline sequence (see **Supp. Section 10.3** for calculating germline distance). **Fig. S3** and **Table S9** report the Spearman’s correlation between model confidence and germline distance for datasets within each developability category. ProGen2 BFD90 had the strongest correlation across all categories (ρ = 0.80), and IgLM perplexities were most correlated to germline for sequences with thermostability, binding, polyreactivity, and immunogenicity labels (**Table S9**). Sequence-only models also had significantly higher correlation to germline than any models that learned over sequence and structure (**Table S10**). BioPython and PyRosetta displayed little to no correlation with germline. Increasing the number of model parameters increased the correlation between model confidence and germline signal (**Fig. S4**). The correlations between model confidence and edit distance to germline were significantly higher than the correlations we reported above between model confidence and fitness value. Additionally, AI-based developability prediction performance was significantly higher for datasets where germline sequences had relatively higher fitness, and lower for datasets where germline sequences had relatively lower fitness (**Fig. S5a)—**a trend not observed for physics-based models (**Fig. S5b**).

We next controlled for germline distance using partial correlation^60^, which measures the relationship between two variables while eliminating influence of a third variable (**Supp. Section 10.4**). Upon eliminating the influence of germline distance, protein language models lost, on average, 40% of their apparent fitness-predictive power, whereas physics-based models were significantly less affected (14%) (**Fig. 6, Table S12**). Our results suggest that the dominance of germline residues in the antibody pretraining data introduces a strong bias toward them in zero-shot prediction. Previous work^61–64^ similarly observed that protein language models capture and exploit phylogenetic distances more readily than biophysical characteristics. The reduction of germline bias we observe with the introduction of structure in the learned representation indicates that this bias can be reduced by expanding the learned representation beyond sequence.

**Figure 6:**
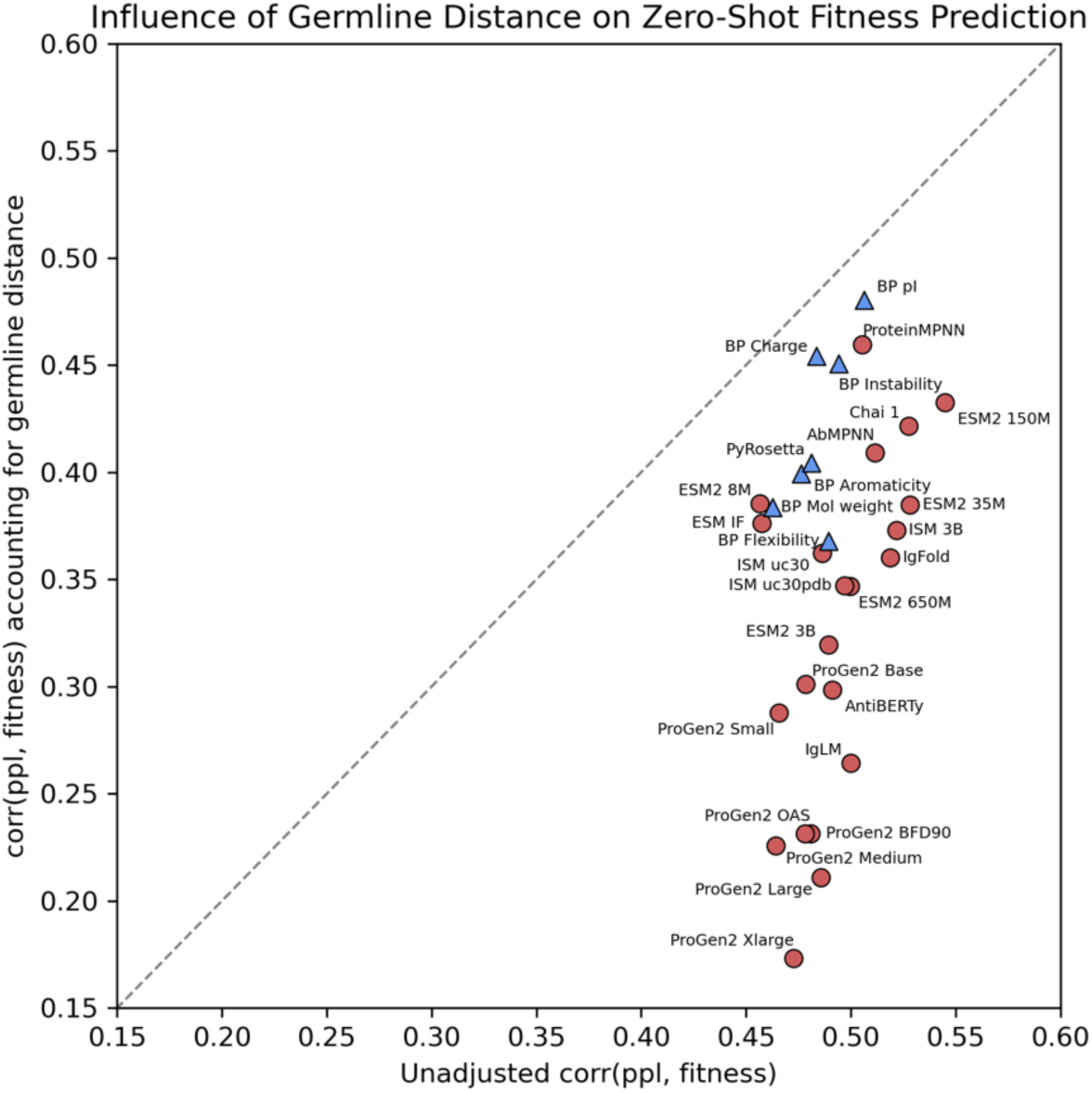
Partial correlation between model confidence and fitness when accounting for germline signal. The *x*-axis shows the correlation between model confidence and fitness, and the *y*-axis presents the partial correlation between model confidence and fitness after accounting for germline signal (see **Supp. Section 10.4**). AI models are displayed as red circles and physics models as blue triangles. To focus on statistically robust results, we include only datasets for which at least five models show statistically significant correlations and for which the mean Spearman correlation across models is at least 0.3 (see datasets in **Table S11**).

## 7 Few-shot results

In the zero-shot setting, we evaluated model uncertainty (perplexity) as a proxy for developability. However, this metric may not fully capture the knowledge encoded during pretraining. To test whether models implicitly learned developability-related features, we used their latent embeddings in a supervised few-shot regression framework. The datasets that satisfy the cutoff of five statistically significant dataset-models and are used in this section are listed in **Table S13**. Results for pharmacokinetics and polyreactivity are omitted due to a lack of at least five statistically significant dataset-model correlations and big enough training datasets. The distribution of Spearman’s correlation coefficients for each fine-tuned model is displayed in **Fig. 7**, and the count of statistically significant correlations per model is displayed in **Fig. S6**.

**Figure 7:**
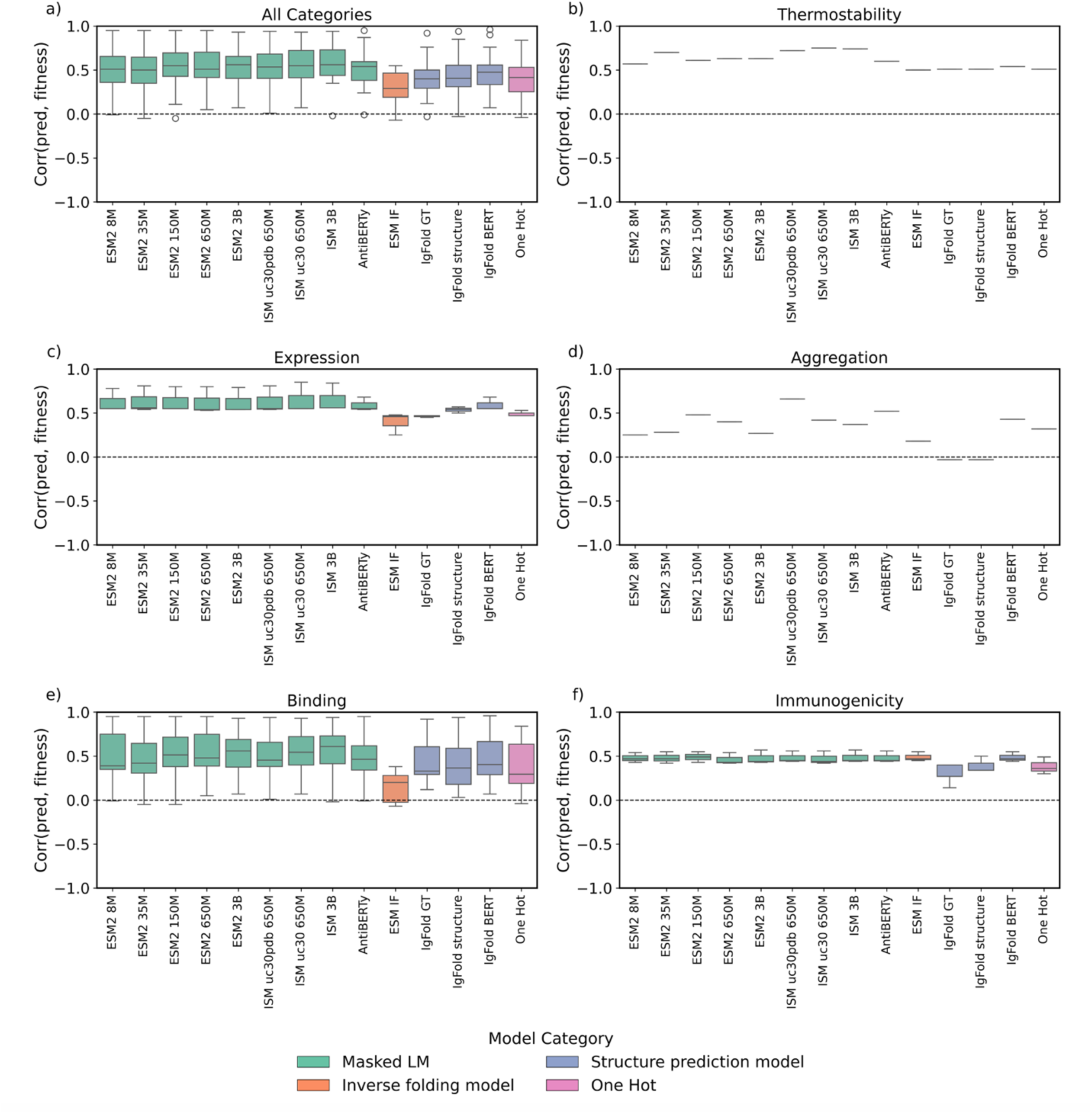
Distribution of few-shot prediction performances for each model. Models are colored based on their architecture, and the *y*-axis displays the range of Spearman’s correlations between predicted developability and true developability label, where a correlation approaching 1.0 is ideal. We only report Spearman’s correlations with datasets that exhibit at least five significant dataset-model p-values to ensure our conclusions are statistically sound.

### 7.1 Few-shot models yield fewer significant correlations than zero-shot

As in the zero-shot setting, we first investigate how many statistically significant Spearman’s correlations each few-shot model has to each developability dataset. The masked language model ESM2 8M displayed the highest count of statistically significant correlations, with 30 correlations or 13% of the total developability datasets (**Fig. S6a**). In order of decreasing count, protein AI models on average displayed significant correlations to 73% of immunogenicity datasets (**Fig. S6h**), 46% of expression datasets (**Fig. S6c**), 9% of binding datasets (**Fig. 6e**), 4% of aggregation datasets (**Fig. S6d**), 6% of thermostability datasets (**Fig. S6b**), 5% of polyreactivity datasets (**Fig. S6g**), and 2% of pharmacokinetic datasets (**Fig. S6f**). This trend is similar to the zero-shot setting, where immunogenicity and expression had the highest count of significant correlations compared to the other developability categories. The overall lower count of significant correlations is in part due to the small size of the developability datasets – most datasets are less than 100 datapoints (**Fig. 2c**) leading to small train/val/test splits and poor performance. To enable a more equitable comparison of model performance, we identified the subset of the datasets that exhibit statistically significant correlations across at least five models (**Table S13**). The corresponding distributions of Spearman’s correlation coefficients for each model with the selected datasets are shown in **Fig. 7**.

### 7.2 Few-shot learning improves performance in all developability categories

Within each developability category, we investigate the average performance of MLPs trained on embeddings from each protein AI model, identify the best performing model, and compare it to the top performing zero-shot model. Thermostability was the least challenging to predict (ρ = 0.61), while aggregation was the most difficult (ρ = 0.32) (**Table S14**). Like the zero-shot setting, expression and immunogenicity showed statistically significant correlations across the largest fraction of models (**Fig. S6**). From the easiest to hardest, the average developability prediction performance across all models was: thermostability > expression > immunogenicity > binding > aggregation.

Embeddings from two protein AI models tied for best performer across all developability properties (ρ = 0.56): ESM2 3B and ISM 3B (**Fig. 7a**, **Table S15**). Most developability categories had a unique top-performing model (**Fig. 7b-g**, **Table S15**). Embeddings from the structure-informed masked language model ISM uc30 650M best predicted thermostability (ρ = 0.75) (**Fig. 7b**), the masked language models ESM2 35M and ISM 3B best predicted expression (ρ = 0.56) (**Fig. 7c**), ISM uc30pdb 650M best predicted aggregation (ρ = 0.66) (**Fig. 7d**), ISM 3B best predicted binding (ρ = 0.61) (**Fig. 7e**), and ESM2 150M best predicted immunogenicity (ρ = 0.49) (**Fig. 7h**, **Table S15**). High performance in one category did not generalize to others. For instance, while the IgFold structure embeddings model was among the best predictors for expression (ρ = 0.54), it was a poor predictor of aggregation (ρ = -0.03). Across all categories, the best few-shot models perform better than the best zero-shot models. In both the zero-shot and few-shot setting, aggregation was among the most challenging. However, there was a change in the other categories, as thermostability became the easiest to predict after finetuning, over expression (**Table S15**).

### 7.3 Architecture impacts few-shot more than zero-shot performance

We next ask whether model architecture influences few-shot performance. Across each developability category, embeddings from masked language models (ρ = 0.55) performed the best (**Table S16**). One hot encoding models performed similarly to inverse folding models and structure predictors (**Table S16**). These results are different to the zero-shot setting, where we found that architecture did not significantly impact performance.

### 7.4 Learned representation does not significantly impact few-shot performance

We next asked whether learned protein representation influences few-shot performance. Embeddings from structure-informed masked language models generally performed best (**Table S17**). Across all developability properties, both ESM2 3B (sequence-only) and ISM 3B (structure-informed) performed equally (ρ = 0.56). ISM uc30pdb 650M (ρ = 0.54) and ESM2 650M (ρ = 0.51) also performed similarly. Some models, such as IgFold, produce multiple learned representations throughout their pipeline. From IgFold, we extract sequence-only AntiBERTy embeddings, structure-informed embeddings after the graph transformer, and further structure-informed embeddings following template incorporation. Performance varied across these representations: AntiBERTy embeddings performed best across all properties (ρ = 0.48), followed by IPA (ρ = 0.41) and the graph transformer (ρ = 0.40). Whereas in the zero-shot setting adding structure improved performance, fine-tuning sequence-only embeddings were sufficient to obtain competitive developability prediction performance (**Table S15**).

### 7.5 General protein embeddings outperform antibody-specific embeddings in few-shot setting

We next assessed how embeddings from models trained specifically on antibody sequences compare to embeddings from general protein models. Embeddings from general protein models dominated at predicting each developability property (**Table S18**). These results differ from the zero-shot setting, where there was more variation in what general protein and antibody-specific models were effective at predicting.

### 7.6 One-hot encodings perform comparably to billion-parameter models in few-shot prediction

We next asked whether the size of the model influences few-shot prediction performance. **Fig. 8** shows that increasing model size did not significantly improve few-shot performance across all categories. For thermostability (**Fig. 8b**), expression (**Fig. 8c**), aggregation (**Fig. 8d**), binding (**Fig. 8e**), and immunogenicity (**Fig. 8f**), one hot encoding performed equally to million and billion parameter models, indicating that simple models given sufficient developability data can approach the accuracy of large pretrained protein models. Developability prediction does not require billion parameter models, neither in zero-shot nor in few-shot.

**Figure 8:**
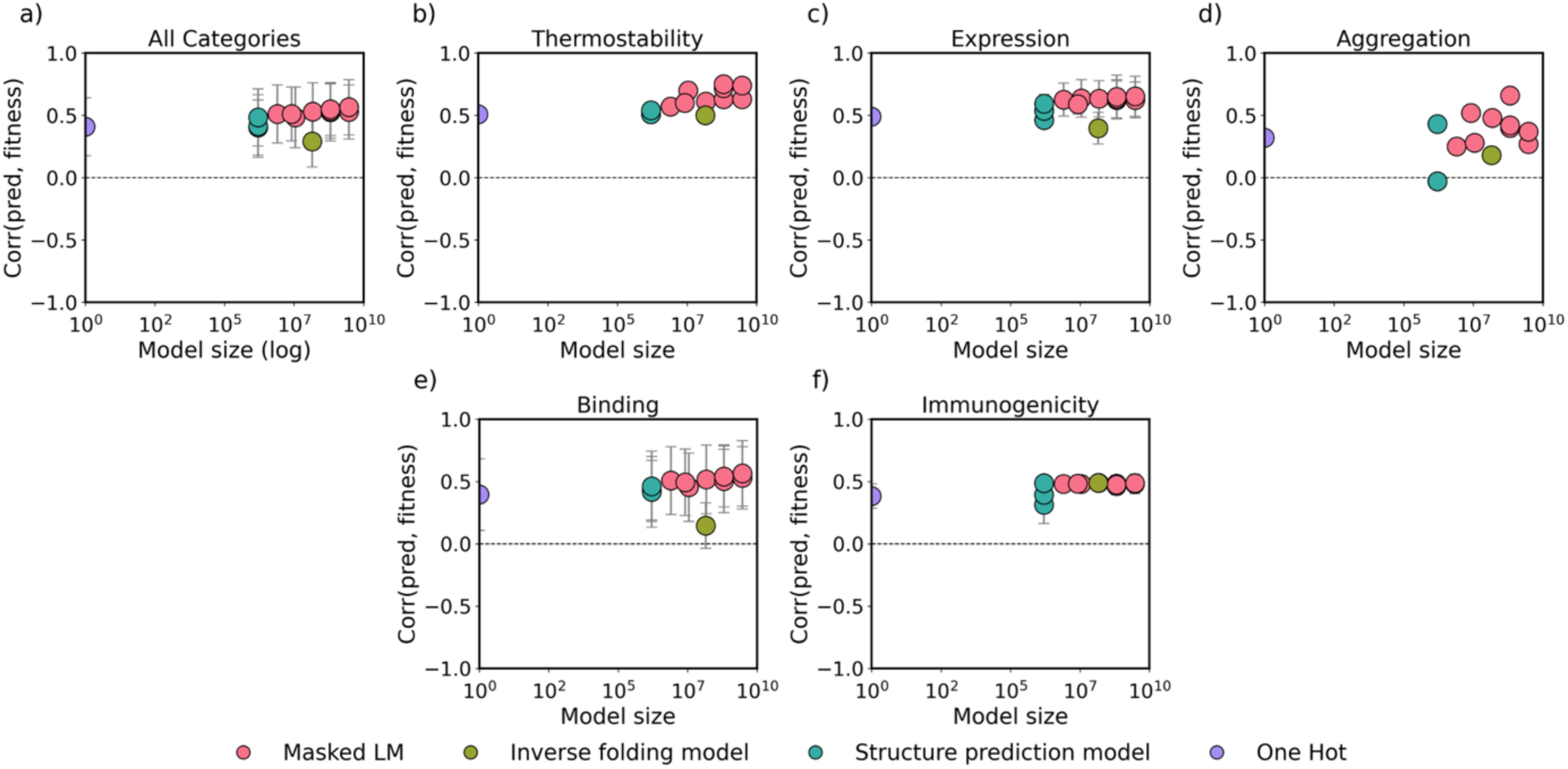
Distribution of few-shot prediction performances for each model compared to parameter size. Models are colored based on their architecture, and the *y*-axis displays the range of Spearman’s correlations between predicted developability and true developability label, where a correlation approaching 1.0 is ideal. We only report Spearman’s correlations with datasets that exhibit at least five significant dataset-model p-values to ensure our conclusions are statistically sound.

### 7.7 Few-shot models lose their germline bias compared to zero-shot models

We next investigated whether models trained on embeddings of pretrained protein AI models are correlated to germline signals, rather than the actual developability label. We calculated the Spearman’s correlation between model’s predicted developability and the Levenstein distance of the antibody from its respective germline sequence (see **Supp. Section 10.3** for calculating germline distance). **Fig. S7** and **Table S19** report the Spearman’s correlation between predicted developability and germline distance. Across all developability properties, the correlation to germline varied significantly, with some exhibiting an inverse correlation and others showing a moderate correlation of ρ < 0.3 (**Table S19**). The germline bias shows no sign of increasing or decreasing with parameter size (**Fig. S8**). Compared to the zero shot models presented earlier, the germline correlations exhibited by few shot models were significantly less.

In the zero-shot setting described earlier, all pretrained protein AI models exhibited a clear germline bias, reflecting their reliance on evolutionary signals encoded during pretraining. In contrast, this bias substantially diminishes in the few-shot setting. Here, model predictions show little to no correlation with germline distance, suggesting that the models are no longer depending on evolutionary similarity to infer developability. Instead, they appear to have learned features more directly related to the experimental labels, resulting in improved predictive performance. The only developability property that consistently retained a notable correlation with germline distance was immunogenicity. This finding aligns with prior reports^65,66^ that antibodies diverging further from their germline sequences tend to have higher immunogenic potential, and so germline distance is inherently correlated with immunogenicity.

## 8 Conclusion

This work benchmarked the zero-shot and few-shot performance of diverse protein AI models in predicting antibody developability properties. We highlighted the models that performed best in each developability category; evaluate how model architecture, learned protein representation, pretraining data composition, and parameter size influence performance; and identified what properties of therapeutic antibodies protein AI models learn.

Based on these observations, we suggest a set of recommendations to consider when developing zero-shot and few-shot antibody developability oracles:

1. Developability properties should be examined separately, as models that perform well in one fitness task may not perform well in any of the others.
2. Incorporating protein structure into the learned representation improves zero-shot and few-shot performance and reduces germline bias.
3. Antibody-specific models generally perform better for binding affinity and immunogenicity prediction, while general protein models perform better at thermostability and expression prediction.
4. Model size does not significantly impact zero-shot prediction.
5. When presented with little fine-tuning data (< 300 sequences), train an MLP on embeddings from a large, billion parameter model. When presented with a lot of fine-tuning data (> 1000 sequences), training an MLP on one-hot encodings may suffice.

The task of generalizable zero-shot and few-shot developability prediction remains unsolved. The protein AI models we evaluated rely on simplified heuristics—distance from germline and evolutionary patterns—rather than capturing the biophysical mechanisms underlying antibody developability. These heuristics do not generalize across diverse developability property landscapes, reflecting an incomplete internal representation. These models learn statistical patterns observed in the training data rather than the physical laws governing folding or function. Recognizing these limitations highlights the need to design models that can integrate richer sources of information to improve generalization.

The current benchmark has several constraints that limit broad conclusions. Not all model architectures are equally represented, with fewer structure-informed masked language models and a predominance of general protein models over antibody-specific models. Additionally, some datasets are small (2 - 20 sequences), resulting in weak statistical power and potential spurious correlations. To ensure robustness, we chose to interpret our results by looking only at datasets that had at least five significant dataset-model correlations from our set of 30 models and 240 datasets. Ideally, all correlations between models and datasets would be significant for a robust comparison to draw conclusions, but this is not the case for a database composed of datasets with varying size and biological assays with varying noise in their readouts. The next iteration of an antibody fitness benchmark could extend FLIP^27^ by evaluating model generalization across diverse mutational regimes—including single, double, and triple mutants—to capture nonlinear epistatic effects^67^. Investigating models trained on low-fitness variants and tested on high-fitness regions could further assess their capacity to guide directed evolution. Comparing machine learning–designed sequences with experimentally evolved variants would provide insight into whether a protein AI model interprets each differently. Pairwise variant ranking frameworks like that used in AbRank^34^ could enhance robustness to assay noise and distribution shifts. Finally, exploring homolog density in training data^58^ and adopting fine-tuning strategies like PLMFit^68^, domain adaptation^69^, and calibration^70^ could improve zero-shot and few-shot prediction. Future improvements to developability oracles may involve integrating developability-labeled data^71^, experimental feedback^72^, in-silico developability data^73^, and physics-based information^74^ to enhance model generalization. Incorporating antigen or environmental context, along with domain adaptation strategies, may help account for covariate shifts^75^ across heterogeneous datasets. Mixing representations, such as combining language model embeddings with one-hot encodings^76^, has shown promise for improving predictive performance, and should be further explored.

Improvements to both benchmarking and developability oracles requires more developability data, as we have found that sequence and structure alone cannot capture all the fitness landscapes of therapeutic antibodies. This ideal developability dataset includes (1) diversity in antibody sequence space with many clonotypes and many types of mutations like substitution and indels; (2) diversity in developability space by including antibodies with good or poor developability properties, as we have limited data of antibodies with poor developability; (3) antibodies with low or high edit distances from germline; and (4) data replicated in multiple labs or environments to estimate the effects of covariates in combining datasets. Hummer et al.^77^ estimated at least 90K computational ΔΔG values must be obtained to train an accurate binding affinity oracle. Properties that we found challenging to predict in our benchmark (immunogenicity, aggregation, pharmacokinetics) may require more data than this estimate, and properties we found easier to predict (expression) may require less data. Although FLAb2 contains millions of datapoints for antibody binding affinity, most are for the same 2 – 3 antigen sequences, making it difficult for us to reproduce the results of Hummer et al. which trained on a more diverse set of antibody-antigen complexes across the entire PDB. Expanding the benchmark with larger, more balanced datasets in the bottom right regime of **Fig. 2 c)** will improve the robustness and interpretability of observed trends.

Open data initiatives such as FLAb2 can accelerate the development of robust computational oracles, provide reproducible benchmarks, and ultimately guide the design of therapeutic antibodies with improved developability. However, more publicly available antibody developability data than what is currently available is necessary to benchmark and train the developability oracles of tomorrow. The continual release of antibody developability data will lead to major improvements in performance of antibody design and screening by leveraging the ingenuity of the global antibody engineering community.

## 9 Acknowledgments

We thank all the experimentalists who created the developability data we collected in FLAb2, as it provides the foundation for benchmarking and developing AI-based therapeutic antibody oracles. We thank Luis Garbinski at GlaxoSmithKline for providing in-house antibody data on binding, expression, and thermostability (labelled as Garbinski et al. unpublished (2023)). We thank Timothy Whitehead for providing MAGMA-seq binding data from Kirby et al^78^. This project was supported by the National Institutes of Health (NIH) grant number R35 GM141881 (JJG) and the National Science Foundation (NSF) Graduate Research Fellowship Program (MC).

## 10 Supplementary

### 10.0 Data availability

Data, scripts, and analyses are available at https://github.com/Graylab/FLAb/releases/tag/v1.0.0.

The datasets collected and used in our analyses are covered by a diverse set of licenses, and we only share datasets in our GitHub that allow redistribution (Attribution 4.0 International, Attribution-NonCommercial 4.0 International, Attribution-NonCommercial-NoDerivatives 4.0 International, MIT, Clear BSD). The datasets not redistributed in the FLAb2 GitHub are from Ginkgo Datapoints^12^ (https://huggingface.co/datasets/ginkgo-datapoints/GDPa1) and NaturalAntibody^35,79^ (https://naturalantibody.com/ab-design/, https://naturalantibody.com/therapeutic-antibody-database/).

**Table S1:**
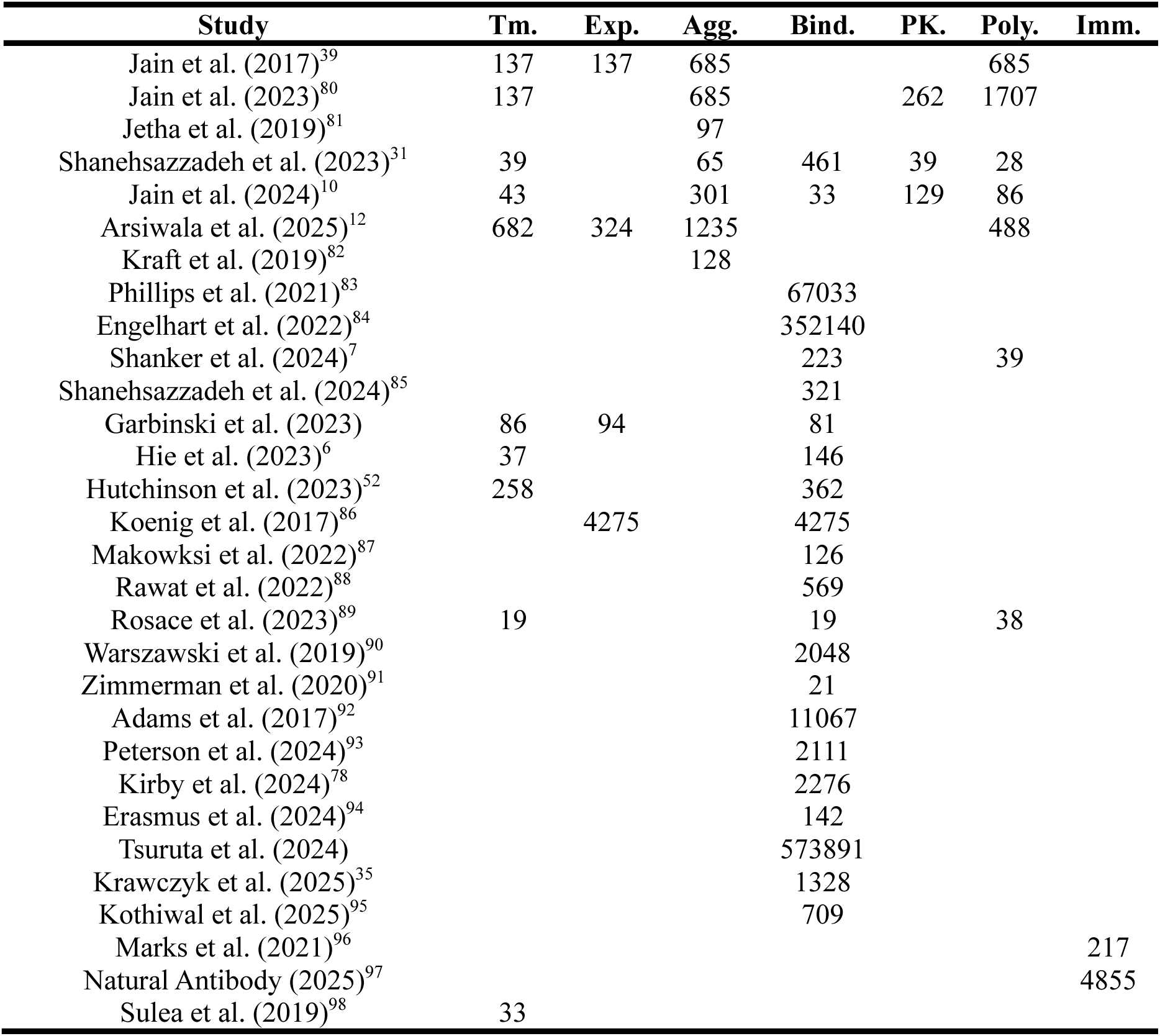
Number of unique antibody sequence-fitness label pairs from antibody datasets. The data provided by Garbinski et al. (2023) is not associated with a publication.

| Study | Tm. | Exp. | Agg. | Bind. | PK. | Poly. | Imm. |
| --- | --- | --- | --- | --- | --- | --- | --- |
| Jain et al. (2017)<sup>39</sup> | 137 | 137 | 685 |  |  | 685 |  |
| Jain et al. (2023)<sup>80</sup> | 137 |  | 685 |  | 262 | 1707 |  |
| Jetha et al. (2019)<sup>81</sup> |  |  | 97 |  |  |  |  |
| Shanehsazzadeh et al. (2023)<sup>31</sup> | 39 |  | 65 | 461 | 39 | 28 |  |
| Jain et al. (2024)<sup>10</sup> | 43 |  | 301 | 33 | 129 | 86 |  |
| Arsiwala et al. (2025)<sup>12</sup> | 682 | 324 | 1235 |  |  | 488 |  |
| Kraft et al. (2019)<sup>82</sup> |  |  | 128 |  |  |  |  |
| Phillips et al. (2021)<sup>83</sup> |  |  |  | 67033 |  |  |  |
| Engelhart et al. (2022)<sup>84</sup> |  |  |  | 352140 |  |  |  |
| Shanker et al. (2024)<sup>7</sup> |  |  |  | 223 |  | 39 |  |
| Shanehsazzadeh et al. (2024)<sup>85</sup> |  |  |  | 321 |  |  |  |
| Garbinski et al. (2023) | 86 | 94 |  | 81 |  |  |  |
| Hie et al. (2023)<sup>6</sup> | 37 |  |  | 146 |  |  |  |
| Hutchinson et al. (2023)<sup>52</sup> | 258 |  |  | 362 |  |  |  |
| Koenig et al. (2017)<sup>86</sup> |  | 4275 |  | 4275 |  |  |  |
| Makowski et al. (2022)<sup>87</sup> |  |  |  | 126 |  |  |  |
| Rawat et al. (2022)<sup>88</sup> |  |  |  | 569 |  |  |  |
| Rosace et al. (2023)<sup>89</sup> | 19 |  |  | 19 |  | 38 |  |
| Warszawski et al. (2019)<sup>90</sup> |  |  |  | 2048 |  |  |  |
| Zimmerman et al. (2020)<sup>91</sup> |  |  |  | 21 |  |  |  |
| Adams et al. (2017)<sup>92</sup> |  |  |  | 11067 |  |  |  |
| Peterson et al. (2024)<sup>93</sup> |  |  |  | 2111 |  |  |  |
| Kirby et al. (2024)<sup>78</sup> |  |  |  | 2276 |  |  |  |
| Erasmus et al. (2024)<sup>94</sup> |  |  |  | 142 |  |  |  |
| Tsuruta et al. (2024) |  |  |  | 573891 |  |  |  |
| Krawczyk et al. (2025)<sup>35</sup> |  |  |  | 1328 |  |  |  |
| Kothiwal et al. (2025)<sup>95</sup> |  |  |  | 709 |  |  |  |
| Marks et al. (2021)<sup>96</sup> |  |  |  |  |  |  | 217 |
| Natural Antibody (2025)<sup>97</sup> |  |  |  |  |  |  | 4855 |
| Sulea et al. (2019)<sup>98</sup> | 33 |  |  |  |  |  |  |

**Figure S1:**
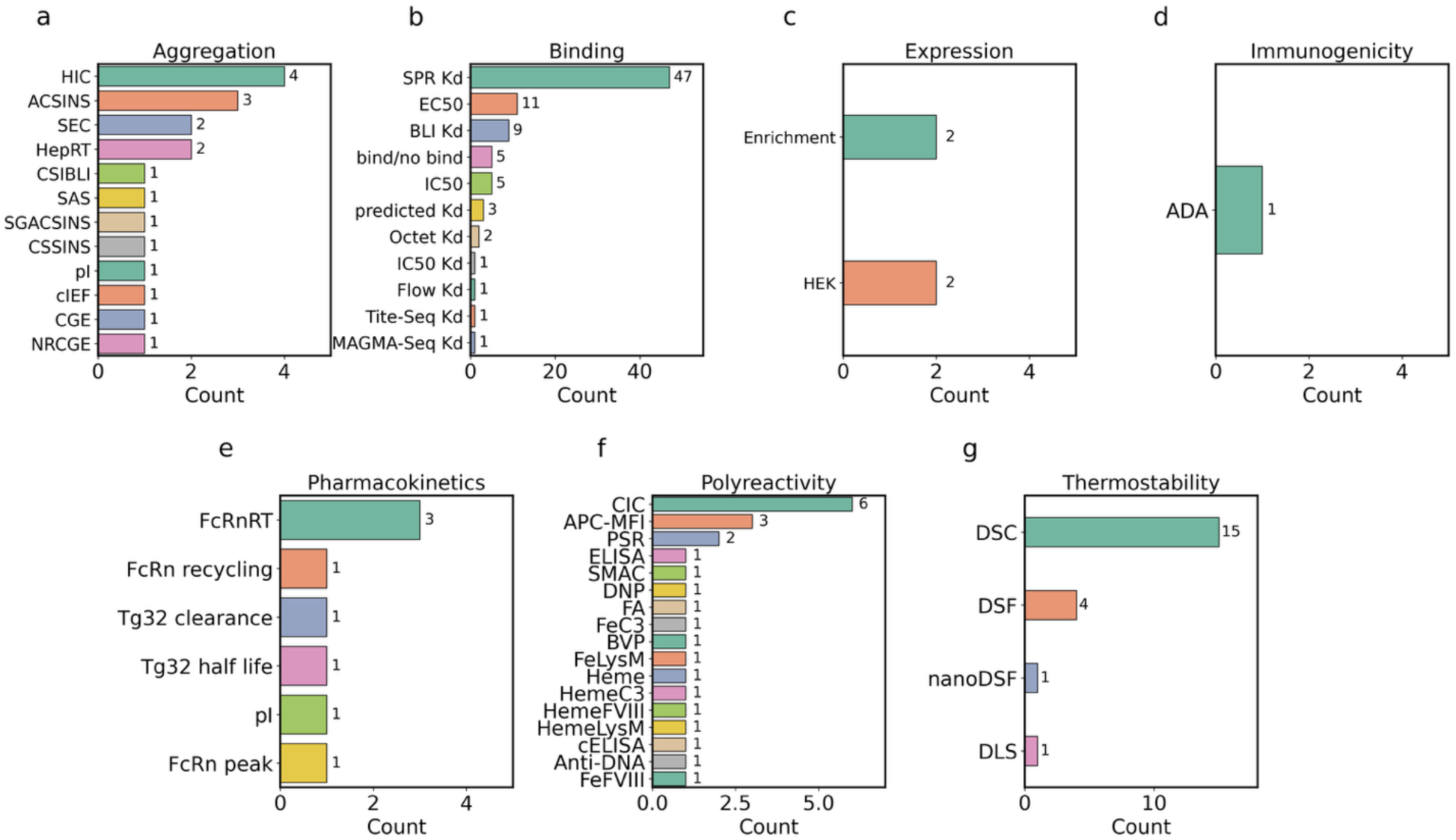
Assays present in each developability category. Each developability property can be measured with a variety of different assays, and we present the counts of each collected assay.

### 10.1 Data composition

### 10.2 Descriptions of developability properties

**Thermostability** ensures an antibody will maintain its structure and function when exposed to heat, particularly during manufacturing, storage, and administration. Antibodies with high thermostability are more likely to remain potent over extended periods and under different storage conditions. Differential scanning fluorimetry (DSF) and calorimetry (DSC) is provided for clinical stage and germline antibodies, with some datasets including point mutations of a starting antibody candidate. The datasets range in size - some are a few hundred antibody variants from the same wildtype^6,10,12,31,52,80,89^, some are collections of diverse antibodies from other public datasets like NbThermo^99^, and some are only 2-3 sequences in dataset size^6^.

**Expression** ensures the production of antibodies in a host cell system, which is necessary to isolate a molecule for further testing and directly affects production yield and cost of manufacturing. Expression data is collected from clinical stage therapeutics^14,39^ and rounds of enrichment of a starting antibody candidate^86^.

**Aggregation** refers to the process of individual antibodies coming together to form larger assemblies, or aggregates. Aggregation can be problematic as it leads to reduced therapeutic efficacy and potentially harmful immune responses. Aggregation data is collected for both therapeutic^12,39,80^ and germline^10^ antibody datasets, and a breadth of assays for measuring this property (**Fig. S1a**).

**Binding affinity** ensures the prolonged physical contact during an interaction between an antibody and target antigen to initiate a mechanism of action: Impacting an antigen’s ability to block pathways (blocking/antagonist); binding a cell surface receptor to activate some downstream response (agonist); initiation of cellular cytotoxicity, phagocytosis, or apoptosis; or receptor internalization and down modulation^100^. Binding affinity data is collected for therapeutic and germline antibodies, in varying sequence diversity from completely different isotypes to single point deep mutational scans.

**Pharmacokinetics** refers to how the body of an animal interacts with an administered therapeutic. A complete pharmacokinetic profile encompasses absorption, distribution, metabolism, and excretion. The pharmacokinetic data collected measures the clearance of a drug through mice^10^, which provides an indication of the drug’s half-life and therapeutic efficacy per dose.

**Polyreactivity** refers to the promiscuity of antibodies to bind to multiple antigens. Although bispecific and trispecific antibodies do exist^1^ and continue to be explored as powerful clinical drug candidates, polyreactivity is generally undesirable as it indicates off-target and non-specific binding. Polyreactivity data is collected for both therapeutic^12,39^ and germline^10^ antibodies.

**Immunogenicity** refers to the elicitation of an undesirable immune response after administration of a therapeutic antibody, leading to the generation of anti-drug antibodies (ADAs). ADAs can recognize and neutralize therapeutic antibodies, reducing their efficacy and potentially causing adverse effects. Minimizing immunogenicity is important for therapeutic antibodies to maintain their efficacy and safety. Immunogenicity datasets report the percent of the patient population that exhibited an ADA response after therapeutic administration^79,96^. Each therapeutic drug in these datasets are tested in different patient populations, so the reported ADA response rates are not directly comparable across drugs.

### 10.3 Protein AI model descriptions

**Table S2:**
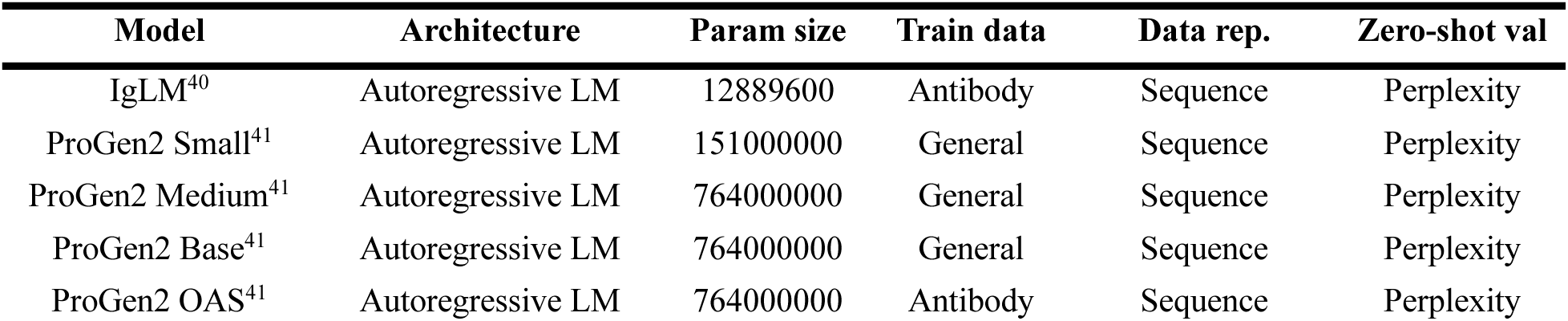
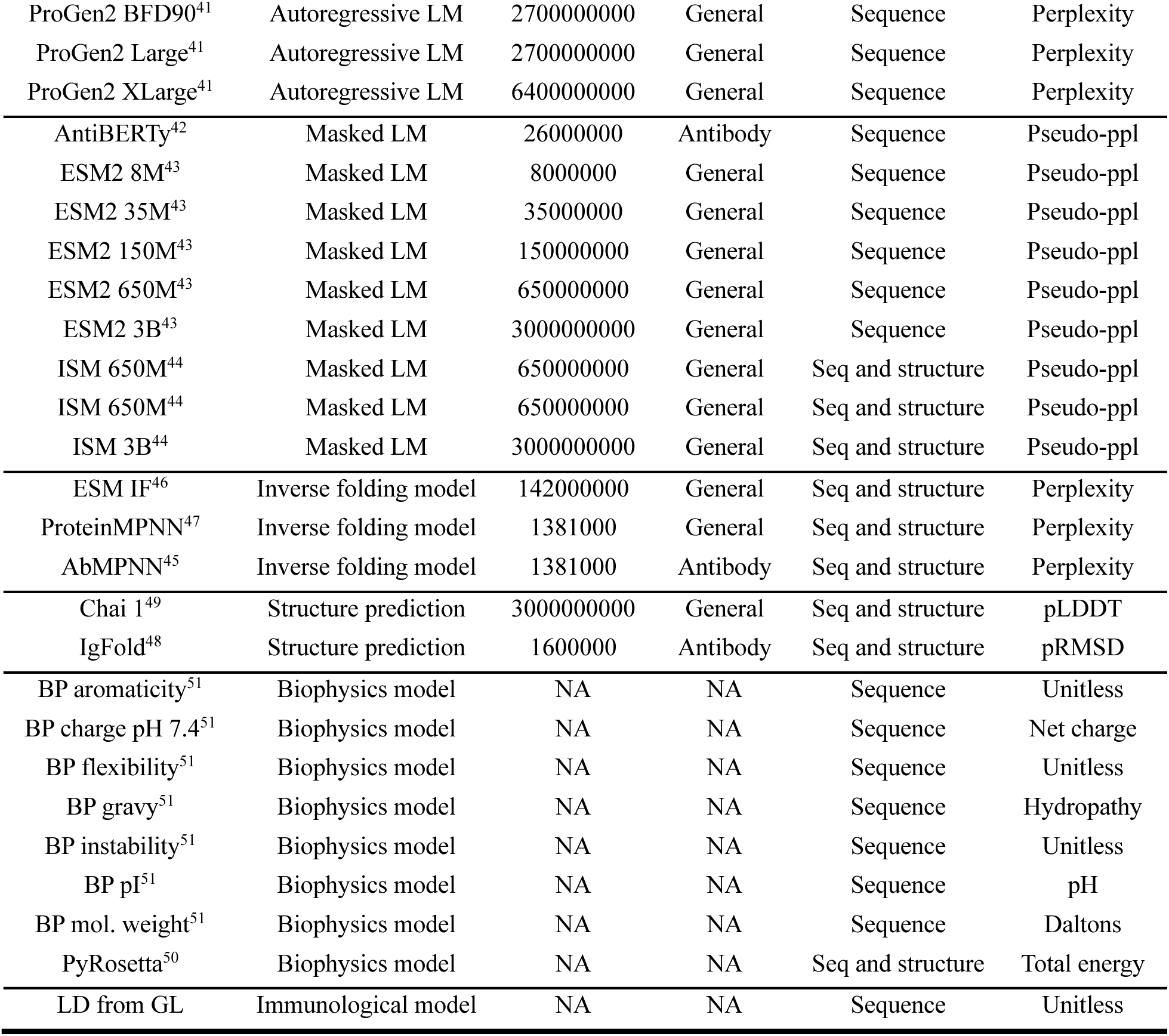
Model descriptions. A description of each benchmarked model is provided in terms of its architecture, the model size, the training data, what representations of protein the model has learned, and the output from the model that is used for zero-shot benchmarking.

#### Decoder-only language models

Decoder-only language models have proven to be effective in generating plausible and novel protein sequences. These models are trained using a next-token prediction objective, where the probability of the next amino acid is influenced by the entire preceding sequence. During training, a database of sequences is utilized to predict *p*(*s*_!_|*s*_“!_), enhancing the model’s ability to generate accurate sequences. We evaluate the zero-shot prediction of therapeutic properties by correlating to the perplexity of each sequence under those models:

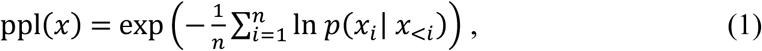

where *x* = (*x*_1_, *x*_2_, … , *x*_3_) is a sequence consisting of *n* tokens.

The decoder models we benchmark are **IgLM** and **ProGen2**. The ProGen2 models come in various sizes, ranging from 151M to 6.4B parameters, pretrained on a mixture of UniRef90 and BFD90 databases. IgLM formulates the design task based on text-infilling using a standard left-to-right decoder (GPT-2), trained on a non-redundant set of 558M antibody sequences obtained from OAS.

#### Encoder-only language models

Encoder-only language models capture comprehensive information in a continuous abstract representation that can be broadly applied. A subset of residues is randomly chosen and replaced with a special mask token. The model is then trained to predict the identities of these masked residues.

In an encoder-only model, an estimation of perplexity can be obtained by calculating the exponential of the negative pseudo-log-likelihood, or pseudo-perplexity:

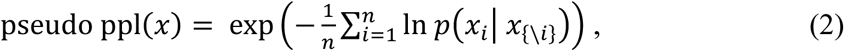

where *x*_{-*i*}_ is the set of all residues except *x_i_*.

In this category of models we focus on **AntiBERTy**, a 26M parameter model pretrained on 558M natural antibody sequences from OAS; **ESM-2**, a suite of models ranging in size from 8M to 3B parameters pretrained on 65M unique sequences from UniRef50 and Uniref90; and **ISM**, which includes the 650M and 3B parameter ESM-2 models distilled with structural representations.

#### Inverse folding models

Generative deep learning architectures that predict protein sequences from structures are known as inverse folding models. For inverse folding models we evaluate the perplexity for each antibody sequence-structure pair:

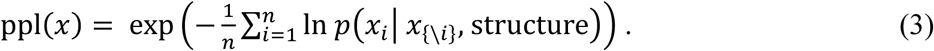

**ESM-IF** uses an autoregressive encoder-decoder architecture, where the model is tasked with recovering the native sequence of the protein from the coordinates of its backbone atoms. **ProteinMPNN** uses a message-passing neural network with 1.4M parameters that predicts protein sequences using the protein backbone geometry. **AbMPNN** uses the same architecture as ProteinMPNN, with weights obtained from training on antibody PDB structures. Structures of all antibody mutants are predicted with Chai-1 prior to scoring with inverse folding models.

#### Structure prediction models

Protein structure prediction models predict the 3D coordinates of a protein backbone given a protein sequence. We measure the network’s structure prediction confidence for each protein sequence as a metric for perplexity, which is pLDDT for **Chai-1**, a general protein structure prediction model, and pRMSD for **IgFold**, an antibody-specific structure prediction model. pLDDT measures per-residue confidence, scaled from 0 to 100 with higher scores indicating higher confidence, whereas pRMSD measures the predicted root-mean squared deviation (RMSD) from the hypothetical true crystal structure, where a value approaching 0 indicates less predicted deviation from the crystal.

#### Physics-based models

We seek to compare the performance of protein AI models versus empirical models of protein energy, which has been a longstanding approach for protein design efforts. **Rosetta**, the classic protein structure prediction and design software, employs an optimized energy function, REF2015, that assesses the energy of atomic interactions within a globular protein. Score functions within Rosetta are composed of weighted sums of various energy terms. Some of these terms correspond to physical forces, such as electrostatics and Van Der Waals interactions, while others represent statistical terms, like the likelihood of observing specific torsion angles in Ramachandran space:

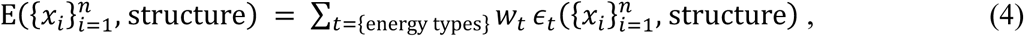

where *∈_i_* is a Rosetta energy term, and *w*_!_is the respective weighted number. Rosetta’s energy calculation does not directly correspond to physical energy units and are instead expressed in Rosetta energy units (REUs). A lower score indicates a higher likelihood of a structure being closer to the native structure. Structures of all antibody mutants are predicted with Chai-1 prior to calculating Rosetta energy.

We use **Biopython** to compute physicochemical properties of antibody sequences: Aromaticity (the frequency of aromatic residues), Charge at pH 7.4 (the net electric charge of the antibody), Flexibility (backbone flexibility based on empirical parameters from amino acid propensities), Gravy (average hydropathy index), Instability index (an estimate of protein stability), and Isoelectric point (pH at which protein carries no net charge).

#### Germline model

Antibodies begin as germline-encoded sequences that diversify through somatic mutation as they mature toward recognizing specific pathogens. We seek to compare the likelihoods generated by the model to the edit distance of the antibody sequence from its respective germline. For each antibody sequence, we retrieve the V gene and J gene calls using ANARCI^92^. The V gene encodes frameworks (FRs) 1-3, complementarity determining regions (CDRs) 1-2, and part of CDR3; and the J gene encodes part of CDR3 and FR4. The edit distance between the antibody sequence and the retrieved sequences from the V and J gene calls are calculated via the Levenstein distance.

### 10.4 Zero-shot results

**Table S3:**
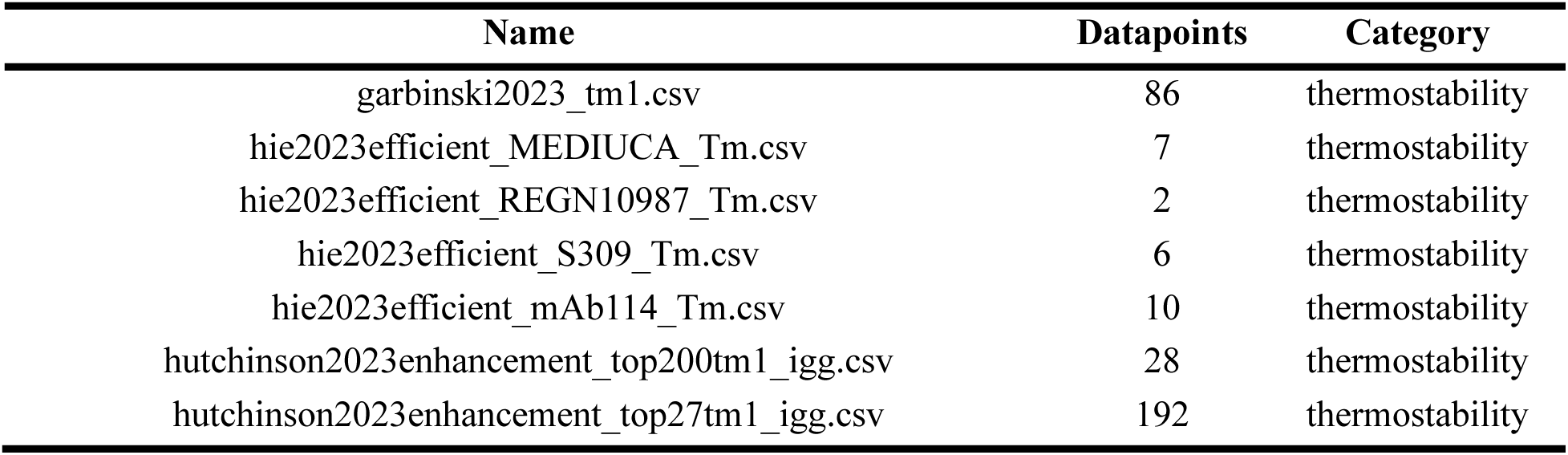
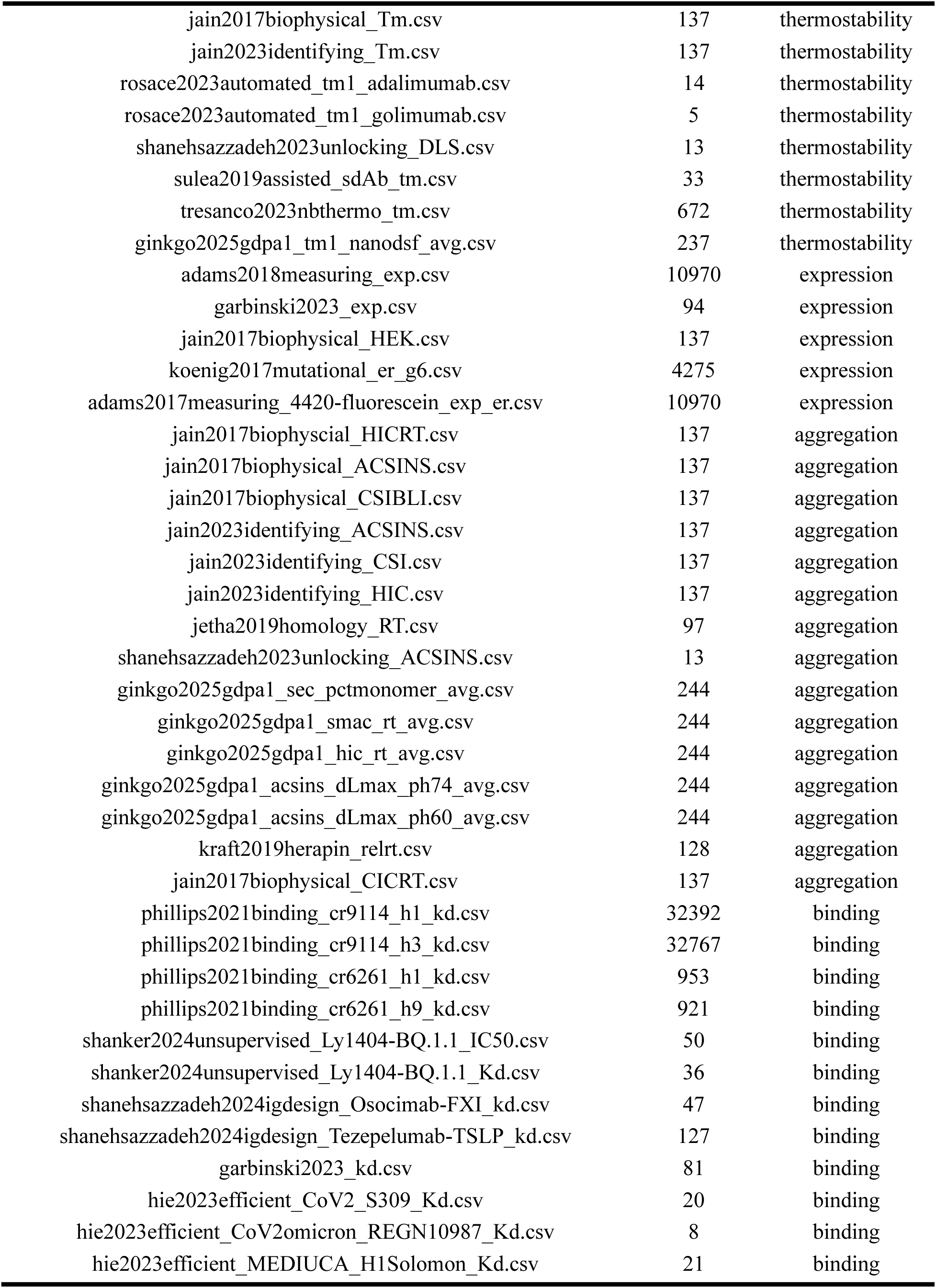
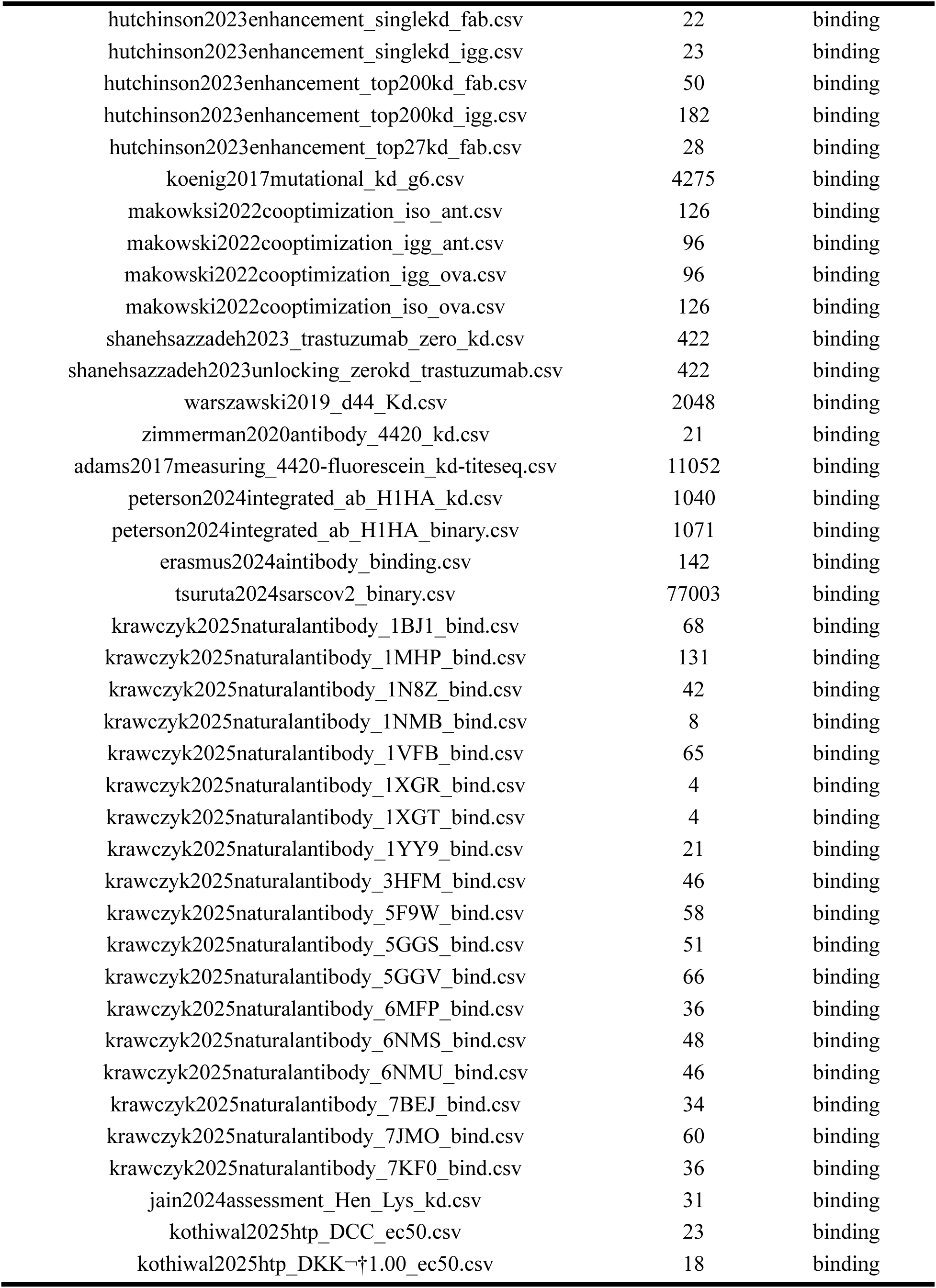
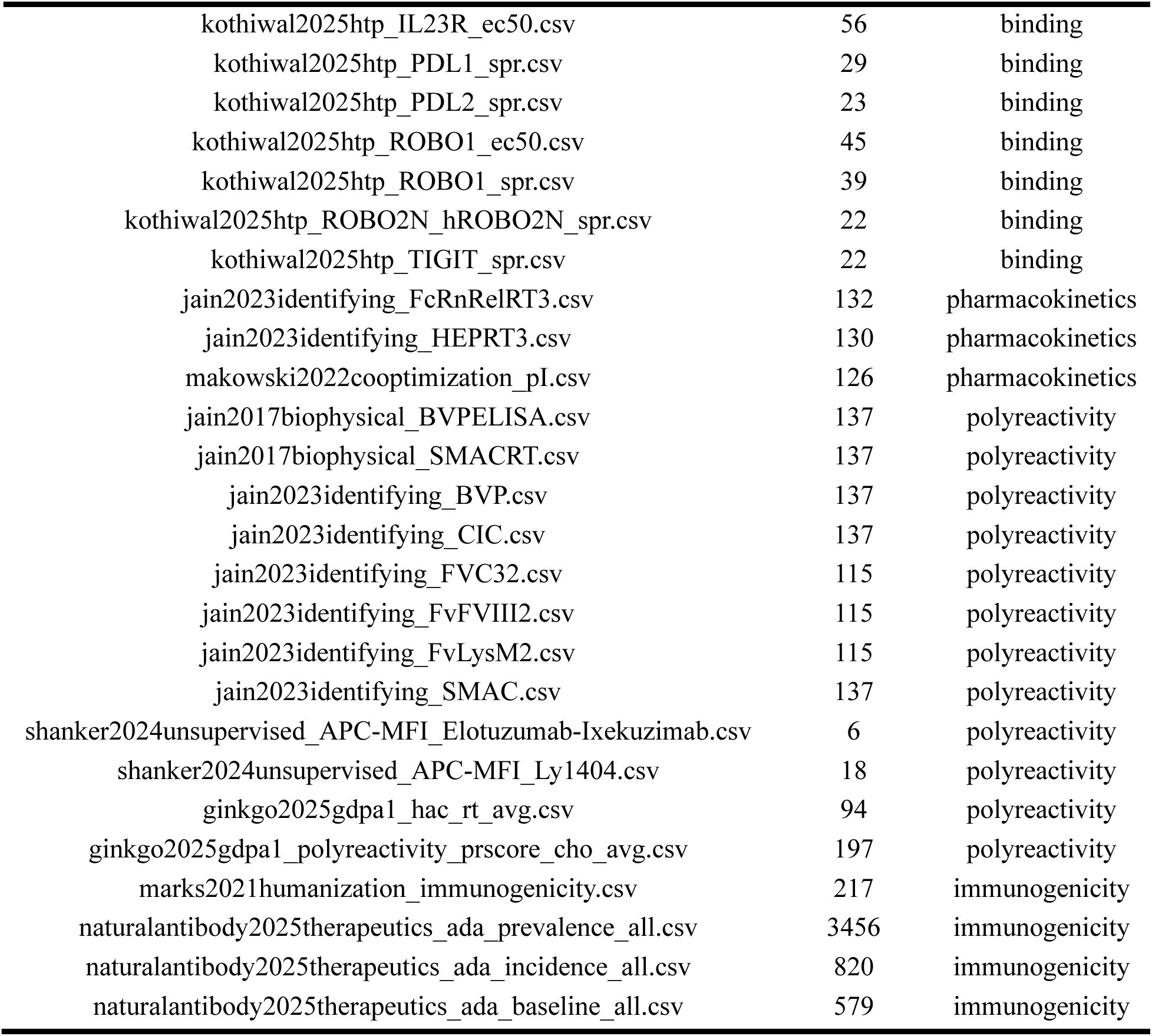
Datasets that pass the 5-significant-model cutoff. Datasets that satisfy the 5-significant-model cutoff and are used for the zero-shot model analyses.

| Name | Datapoints | Category |
| --- | --- | --- |
| garbinski2023_tm1.csv | 86 | thermostability |
| hie2023efficient_MEDIUCA_Tm.csv | 7 | thermostability |
| hie2023efficient_REGN10987_Tm.csv | 2 | thermostability |
| hie2023efficient_S309_Tm.csv | 6 | thermostability |
| hie2023efficient_mAb114_Tm.csv | 10 | thermostability |
| hutchinson2023enhancement_top200tm1_igg.csv | 28 | thermostability |
| hutchinson2023enhancement_top27tm1_igg.csv | 192 | thermostability |
| jain2017biophysical_Tm.csv | 137 | thermostability |
| jain2023identifying_Tm.csv | 137 | thermostability |
| rosace2023automated_tm1_adalimumab.csv | 14 | thermostability |
| rosace2023automated_tm1_golimumab.csv | 5 | thermostability |
| shanehsazzadeh2023unlocking_DLS.csv | 13 | thermostability |
| sulea2019assisted_sdAb_tm.csv | 33 | thermostability |
| tresanco2023nbthermo_tm.csv | 672 | thermostability |
| ginkgo2025gdpa1_tm1_nanodsf_avg.csv | 237 | thermostability |
| adams2018measuring_exp.csv | 10970 | expression |
| garbinski2023_exp.csv | 94 | expression |
| jain2017biophysical_HEK.csv | 137 | expression |
| koenig2017mutational_er_g6.csv | 4275 | expression |
| adams2017measuring_4420-fluorescein_exp_er.csv | 10970 | expression |
| jain2017biophyscial_HICRT.csv | 137 | aggregation |
| jain2017biophysical_ACSINS.csv | 137 | aggregation |
| jain2017biophysical_CSIBLI.csv | 137 | aggregation |
| jain2023identifying_ACSINS.csv | 137 | aggregation |
| jain2023identifying_CSI.csv | 137 | aggregation |
| jain2023identifying_HIC.csv | 137 | aggregation |
| jetha2019homology_RT.csv | 97 | aggregation |
| shanehsazzadeh2023unlocking_ACSINS.csv | 13 | aggregation |
| ginkgo2025gdpa1_sec_pctmonomer_avg.csv | 244 | aggregation |
| ginkgo2025gdpa1_smac_rt_avg.csv | 244 | aggregation |
| ginkgo2025gdpa1_hic_rt_avg.csv | 244 | aggregation |
| ginkgo2025gdpa1_acsins_dLmax_ph74_avg.csv | 244 | aggregation |
| ginkgo2025gdpa1_acsins_dLmax_ph60_avg.csv | 244 | aggregation |
| kraft2019herapin_relrt.csv | 128 | aggregation |
| jain2017biophysical_CICRT.csv | 137 | aggregation |
| phillips2021binding_cr9114_h1_kd.csv | 32392 | binding |
| phillips2021binding_cr9114_h3_kd.csv | 32767 | binding |
| phillips2021binding_cr6261_h1_kd.csv | 953 | binding |
| phillips2021binding_cr6261_h9_kd.csv | 921 | binding |
| shanker2024unsupervised_Ly1404-BQ.1.1_IC50.csv | 50 | binding |
| shanker2024unsupervised_Ly1404-BQ.1.1_Kd.csv | 36 | binding |
| shanehsazzadeh2024igdesign_Osocimab-FXI_kd.csv | 47 | binding |
| shanehsazzadeh2024igdesign_Tezepelumab-TSLP_kd.csv | 127 | binding |
| garbinski2023_kd.csv | 81 | binding |
| hie2023efficient_CoV2_S309_Kd.csv | 20 | binding |
| hie2023efficient_CoV2omicron_REGN10987_Kd.csv | 8 | binding |
| hie2023efficient_MEDIUCA_H1Solomon_Kd.csv | 21 | binding |
| hutchinson2023enhancement_singlekd_fab.csv | 22 | binding |
| hutchinson2023enhancement_singlekd_igg.csv | 23 | binding |
| hutchinson2023enhancement_top200kd_fab.csv | 50 | binding |
| hutchinson2023enhancement_top200kd_igg.csv | 182 | binding |
| hutchinson2023enhancement_top27kd_fab.csv | 28 | binding |
| koenig2017mutational_kd_g6.csv | 4275 | binding |
| makowski2022cooptimization_iso_ant.csv | 126 | binding |
| makowski2022cooptimization_igg_ant.csv | 96 | binding |
| makowski2022cooptimization_igg_ova.csv | 96 | binding |
| makowski2022cooptimization_iso_ova.csv | 126 | binding |
| shanehsazzadeh2023_trastuzumab_zero_kd.csv | 422 | binding |
| shanehsazzadeh2023unlocking_zerokd_trastuzumab.csv | 422 | binding |
| warszawski2019_d44_Kd.csv | 2048 | binding |
| zimmerman2020antibody_4420_kd.csv | 21 | binding |
| adams2017measuring_4420-fluorescein_kd-titeseq.csv | 11052 | binding |
| peterson2024integrated_ab_H1HA_kd.csv | 1040 | binding |
| peterson2024integrated_ab_H1HA_binary.csv | 1071 | binding |
| erasmus2024aintibody_binding.csv | 142 | binding |
| tsuruta2024sarscov2_binary.csv | 77003 | binding |
| krawczyk2025naturalantibody_1BJ1_bind.csv | 68 | binding |
| krawczyk2025naturalantibody_1MHP_bind.csv | 131 | binding |
| krawczyk2025naturalantibody_1N8Z_bind.csv | 42 | binding |
| krawczyk2025naturalantibody_1NMB_bind.csv | 8 | binding |
| krawczyk2025naturalantibody_1VFB_bind.csv | 65 | binding |
| krawczyk2025naturalantibody_1XGR_bind.csv | 4 | binding |
| krawczyk2025naturalantibody_1XGT_bind.csv | 4 | binding |
| krawczyk2025naturalantibody_1YY9_bind.csv | 21 | binding |
| krawczyk2025naturalantibody_3HFM_bind.csv | 46 | binding |
| krawczyk2025naturalantibody_5F9W_bind.csv | 58 | binding |
| krawczyk2025naturalantibody_5GGS_bind.csv | 51 | binding |
| krawczyk2025naturalantibody_5GGV_bind.csv | 66 | binding |
| krawczyk2025naturalantibody_6MFP_bind.csv | 36 | binding |
| krawczyk2025naturalantibody_6NMS_bind.csv | 48 | binding |
| krawczyk2025naturalantibody_6NMU_bind.csv | 46 | binding |
| krawczyk2025naturalantibody_7BEJ_bind.csv | 34 | binding |
| krawczyk2025naturalantibody_7JMO_bind.csv | 60 | binding |
| krawczyk2025naturalantibody_7KF0_bind.csv | 36 | binding |
| jain2024assessment_Hen_Lys_kd.csv | 31 | binding |
| kothiwal2025htp_DCC_ec50.csv | 23 | binding |
| kothiwal2025htp_DKK-†1.00_ec50.csv | 18 | binding |
| kothiwal2025htp_IL23R_ec50.csv | 56 | binding |
| kothiwal2025htp_PDL1_spr.csv | 29 | binding |
| kothiwal2025htp_PDL2_spr.csv | 23 | binding |
| kothiwal2025htp_ROBO1_ec50.csv | 45 | binding |
| kothiwal2025htp_ROBO1_spr.csv | 39 | binding |
| kothiwal2025htp_ROBO2N_hROBO2N_spr.csv | 22 | binding |
| kothiwal2025htp_TIGIT_spr.csv | 22 | binding |
| jain2023identifying_FcRnRelRT3.csv | 132 | pharmacokinetics |
| jain2023identifying_HEPRT3.csv | 130 | pharmacokinetics |
| makowski2022cooptimization_pI.csv | 126 | pharmacokinetics |
| jain2017biophysical_BVPELISA.csv | 137 | polyreactivity |
| jain2017biophysical_SMACRT.csv | 137 | polyreactivity |
| jain2023identifying_BVP.csv | 137 | polyreactivity |
| jain2023identifying_CIC.csv | 137 | polyreactivity |
| jain2023identifying_FVC32.csv | 115 | polyreactivity |
| jain2023identifying_FvFVIII2.csv | 115 | polyreactivity |
| jain2023identifying_FvLysM2.csv | 115 | polyreactivity |
| jain2023identifying_SMAC.csv | 137 | polyreactivity |
| shanker2024unsupervised_APC-MFI_Elotuzumab-Ixekuzimab.csv | 6 | polyreactivity |
| shanker2024unsupervised_APC-MFI_Ly1404.csv | 18 | polyreactivity |
| ginkgo2025gdpa1_hac_rt_avg.csv | 94 | polyreactivity |
| ginkgo2025gdpa1_polyreactivity_prscore_cho_avg.csv | 197 | polyreactivity |
| marks2021humanization_immunogenicity.csv | 217 | immunogenicity |
| naturalantibody2025therapeutics_ada_prevalence_all.csv | 3456 | immunogenicity |
| naturalantibody2025therapeutics_ada_incidence_all.csv | 820 | immunogenicity |
| naturalantibody2025therapeutics_ada_baseline_all.csv | 579 | immunogenicity |

**Table S4:** Zero-shot performance summary across all models. Tabulated correlation between model confidence and actual fitness score, averaged across all models. Some of the assay labels are inverted so that a positive correlation between fitness and assay value always indicates improved model performance.

| All Categories | Tm. | Exp. | Agg. | Bind. | PK. | Poly. | Imm. |
| --- | --- | --- | --- | --- | --- | --- | --- |
| 0.01 | 0.06 | **0.22** | -0.07 | 0.05 | -0.06 | 0.02 | 0.07 |

**Table S5:** Zero-shot performance summary for each individual model. Tabulated correlation between model confidence and actual fitness score, for each model.

| Scoring Method | All Categories | Tm. | Exp. | Agg. | Bind. | PK. | Poly. | Imm. |
| --- | --- | --- | --- | --- | --- | --- | --- | --- |
| IgLM | -0.01 | -0.05 | 0.17 | -0.16 | 0.08 | -0.18 | -0.02 | 0.21 |
| ProGen2 Small | 0.00 | 0.13 | 0.39 | -0.15 | 0.17 | -0.05 | -0.07 | 0.10 |
| ProGen2 Medium | 0.08 | 0.09 | 0.36 | -0.19 | **0.18** | -0.13 | -0.03 | **0.34** |
| ProGen2 Base | -0.02 | 0.02 | 0.38 | -0.15 | 0.01 | -0.16 | -0.02 | 0.16 |
| ProGen2 OAS | 0.04 | 0.09 | 0.12 | -0.18 | 0.11 | -0.18 | -0.03 | 0.12 |
| ProGen2 BFD90 | -0.02 | 0.14 | 0.35 | -0.17 | 0.12 | -0.15 | -0.04 | 0.33 |
| ProGen2 Large | 0.00 | 0.01 | 0.36 | -0.15 | 0.09 | -0.13 | -0.04 | 0.17 |
| ProGen2 XLarge | -0.01 | 0.07 | 0.32 | -0.15 | 0.10 | -0.19 | -0.03 | 0.28 |
| AntiBERTy | 0.00 | -0.14 | 0.24 | -0.11 | 0.06 | -0.16 | 0.00 | 0.17 |
| ESM2 8M | -0.04 | 0.01 | 0.26 | -0.08 | -0.03 | -0.10 | -0.05 | -0.10 |
| ESM2 35M | -0.04 | 0.14 | 0.30 | -0.10 | -0.06 | -0.10 | -0.02 | -0.14 |
| ESM2 150M | -0.04 | 0.00 | **0.44** | -0.12 | 0.01 | -0.15 | -0.07 | -0.03 |
| ESM2 650M | 0.04 | 0.15 | 0.37 | -0.07 | 0.12 | -0.07 | 0.04 | -0.01 |
| ESM2 3B | 0.03 | 0.16 | 0.34 | -0.05 | 0.14 | -0.12 | 0.02 | -0.04 |
| ISM 650M | 0.06 | 0.16 | 0.34 | -0.11 | 0.13 | -0.12 | 0.07 | 0.01 |
| ISM 3B | 0.06 | 0.05 | 0.38 | -0.08 | 0.13 | -0.12 | 0.08 | 0.09 |
| ESM IF | 0.01 | 0.25 | 0.29 | -0.05 | -0.04 | -0.08 | 0.08 | 0.10 |
| ProteinMPNN | 0.04 | -0.05 | 0.09 | -0.07 | 0.04 | -0.08 | 0.09 | 0.08 |
| AbMPNN | 0.00 | -0.08 | 0.14 | -0.11 | 0.06 | -0.12 | 0.13 | 0.07 |
| Chai 1 | 0.01 | **0.45** | 0.38 | -0.17 | 0.01 | -0.12 | 0.04 | 0.11 |
| IgFold | **0.12** | 0.08 | 0.23 | -0.17 | 0.18 | -0.16 | 0.16 | 0.08 |
| PyRosetta | -0.02 | -0.01 | -0.13 | -0.04 | 0.07 | -0.06 | -0.04 | 0.03 |
| BP aromaticity | 0.06 | 0.01 | 0.07 | 0.10 | 0.04 | 0.11 | 0.15 | -0.02 |
| BP charge at pH 7.4 | 0.00 | -0.08 | 0.05 | **0.29** | -0.11 | **0.59** | **0.18** | 0.01 |
| BP flexibility | -0.07 | 0.11 | -0.14 | -0.13 | -0.03 | -0.23 | 0.00 | 0.02 |
| BP gravy | 0.04 | 0.02 | 0.05 | -0.01 | 0.06 | -0.08 | -0.05 | 0.03 |
| BP instability index | -0.06 | 0.04 | 0.07 | 0.13 | -0.15 | 0.15 | 0.00 | -0.14 |
| BP isoelectric point | 0.01 | -0.08 | 0.04 | 0.18 | -0.13 | 0.54 | 0.16 | 0.04 |
| BP molecular weight | 0.02 | -0.04 | 0.11 | 0.08 | -0.06 | 0.06 | 0.06 | -0.08 |
| LD from GL | 0.00 | 0.04 | 0.14 | -0.18 | 0.11 | -0.19 | -0.07 | 0.19 |

**Table S6:** Zero-shot performance across architectures. Autoregressive language models include IgLM and the ProGen2 suite. Masked language models include AntiBERTy, the ESM2 suite, and the ISM suite. Inverse folding models include AbMPNN, ESM IF, and ProteinMPNN. Structure predictors include Chai-1 and IgFold. Sequence-based physics models include BioPython. Structure-based physics models include PyRosetta.

| Architecture | All Categories | Tm. | Exp. | Agg. | Bind. | PK. | Poly. | Imm. |
| --- | --- | --- | --- | --- | --- | --- | --- | --- |
| Autoregressive LMs | 0.01 | 0.06 | 0.31 | -0.16 | **0.11** | -0.15 | -0.03 | **0.21** |
| Masked LMs | 0.01 | 0.07 | **0.33** | -0.09 | 0.06 | -0.12 | 0.01 | -0.01 |
| Inverse folding | 0.02 | 0.04 | 0.17 | -0.08 | 0.02 | -0.09 | **0.10** | 0.08 |
| Structure predictors | **0.07** | **0.27** | 0.30 | -0.17 | 0.09 | -0.14 | **0.10** | 0.10 |
| Seq-based physics | 0.00 | 0.00 | 0.04 | **0.09** | -0.05 | **0.16** | 0.07 | -0.02 |
| Struc-based physics | -0.02 | -0.01 | -0.13 | -0.04 | 0.07 | -0.06 | -0.04 | 0.03 |

**Table S7:** Zero-shot performance across sequence-only and structure-informed models. Masked language modes without structure are AntiBERTY and the ESM2 suite. Masked language models with structure include the ISM suite. Inverse folding models are ESM-IF, ProteinMPNN, and AbMPNN. Structure predictors are Chai-1 and IgFold. Sequence-only masked language and causal language models are the ESM2 suite, AntiBERTY, IgLM, and ProGen2.

| Learned rep. | All Categories | Tm. | Exp. | Agg. | Bind. | PK. | Poly. | Imm. |
| --- | --- | --- | --- | --- | --- | --- | --- | --- |
| MLMs w/o structure | -0.01 | 0.05 | 0.33 | -0.09 | 0.04 | -0.12 | -0.01 | -0.02 |
| MLM w/ structure | 0.06 | 0.11 | **0.36** | -0.10 | **0.13** | -0.12 | 0.08 | 0.05 |
| Inverse folding | 0.02 | 0.04 | 0.17 | **-0.08** | 0.02 | **-0.09** | **0.10** | 0.08 |
| Structure predictor | **0.07** | **0.27** | 0.30 | -0.17 | 0.09 | -0.14 | **0.10** | **0.10** |
| MLMs and CLMs | 0.01 | 0.06 | 0.32 | -0.13 | 0.09 | -0.13 | -0.01 | **0.10** |

**Table S8:** Zero-shot performance across general protein and antibody-specific AI models. General protein AI models include the ProGen2 suite, the ESM2 suite, the ISM suite, ESM IF, ProteinMPNN, and Chai-1. Antibody specific models include IgLM, AntiBERTy, AbMPNN, and IgFold.

| Protein family | All categories | Tm. | Exp. | Agg. | Bind. | PK. | Poly. | Imm. |
| --- | --- | --- | --- | --- | --- | --- | --- | --- |
| General protein | 0.01 | **0.11** | **0.32** | **-0.12** | 0.07 | **-0.12** | 0.00 | 0.09 |
| Antibody specific | **0.03** | -0.05 | 0.19 | -0.14 | **0.09** | -0.16 | **0.07** | **0.13** |

**Figure S2:**
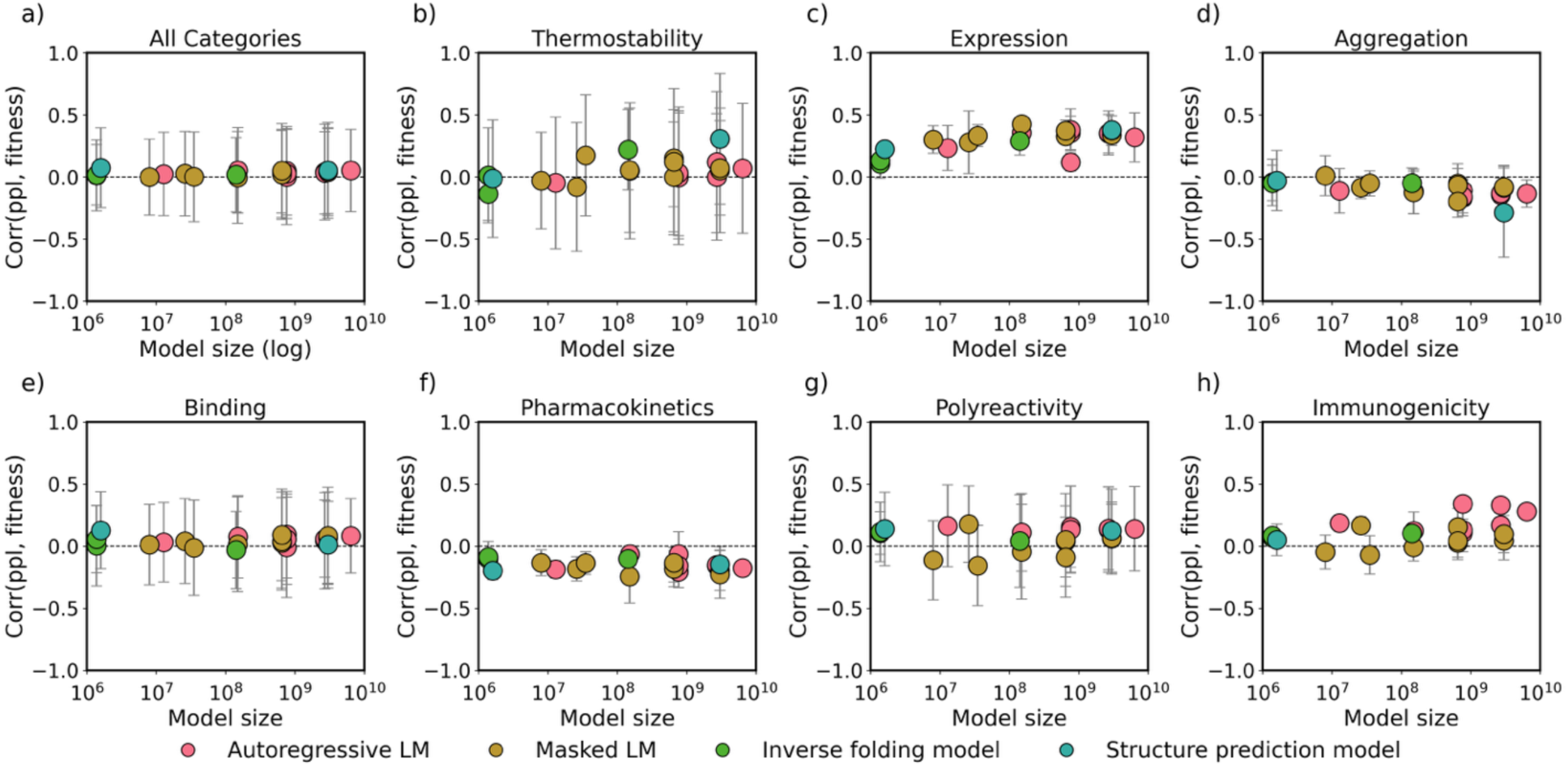
Distribution of zero-shot prediction performances for each model compared to parameter size. Models are colored based on their architecture, and the *y*-axis displays the range of Spearman’s correlations between model confidence and developability label, where a correlation approaching 1.0 is ideal. We only report Spearman’s correlations with datasets that exhibit at least five significant dataset-model p values to ensure our conclusions are statistically sound.

**Figure S3:**
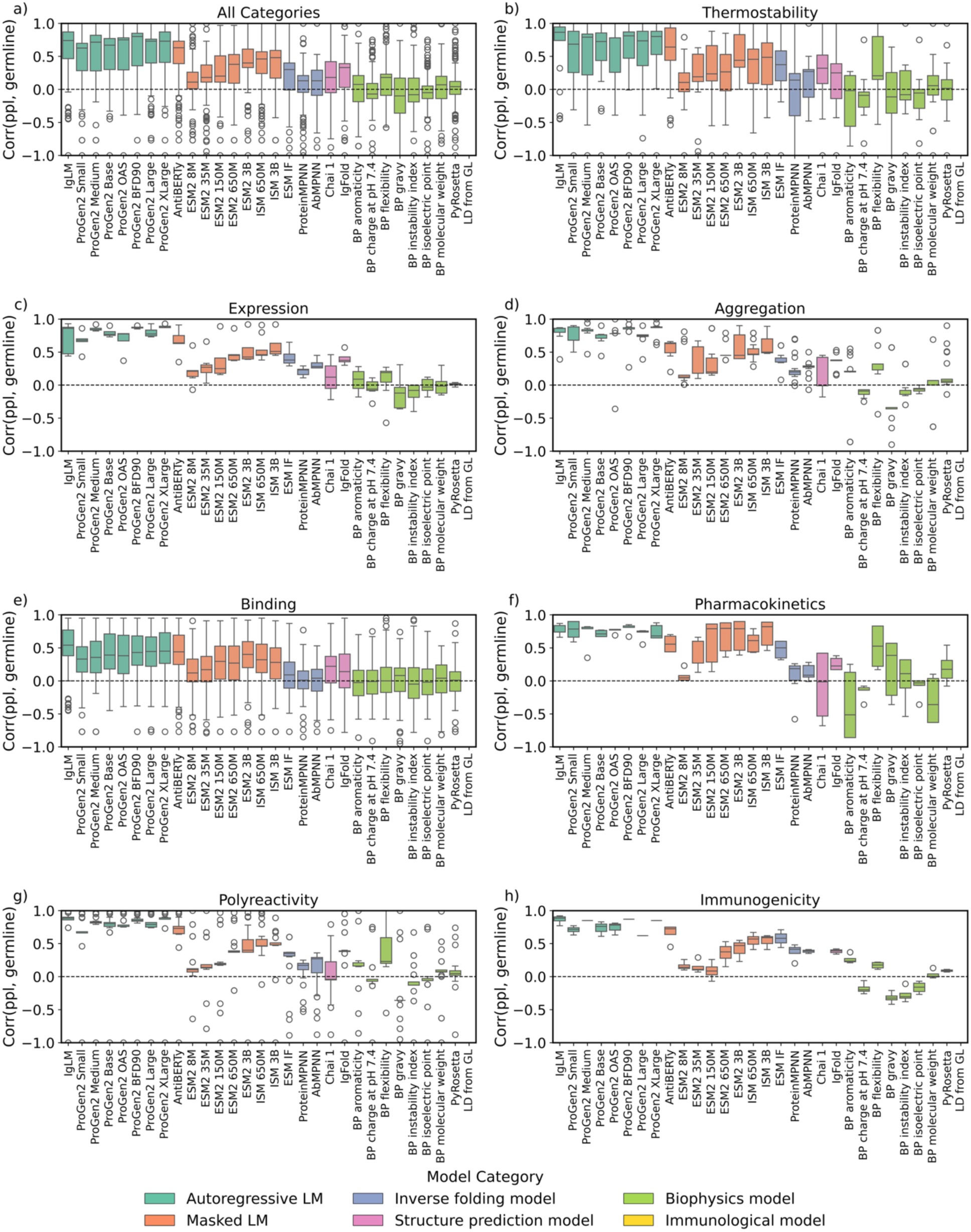
Distribution of zero-shot prediction correlations with germline. Models are colored based on their architecture, and the *y*-axis displays the range of Spearman’s correlations between model confidence and distance from germline. A correlation approaching 1.0 indicates the model is highly confident in germline sequences and highly unconfident in sequences far from germline. We only report Spearman’s correlations with datasets that exhibit at least five significant dataset-model p-values to ensure our conclusions are statistically sound.

**Table S9:**
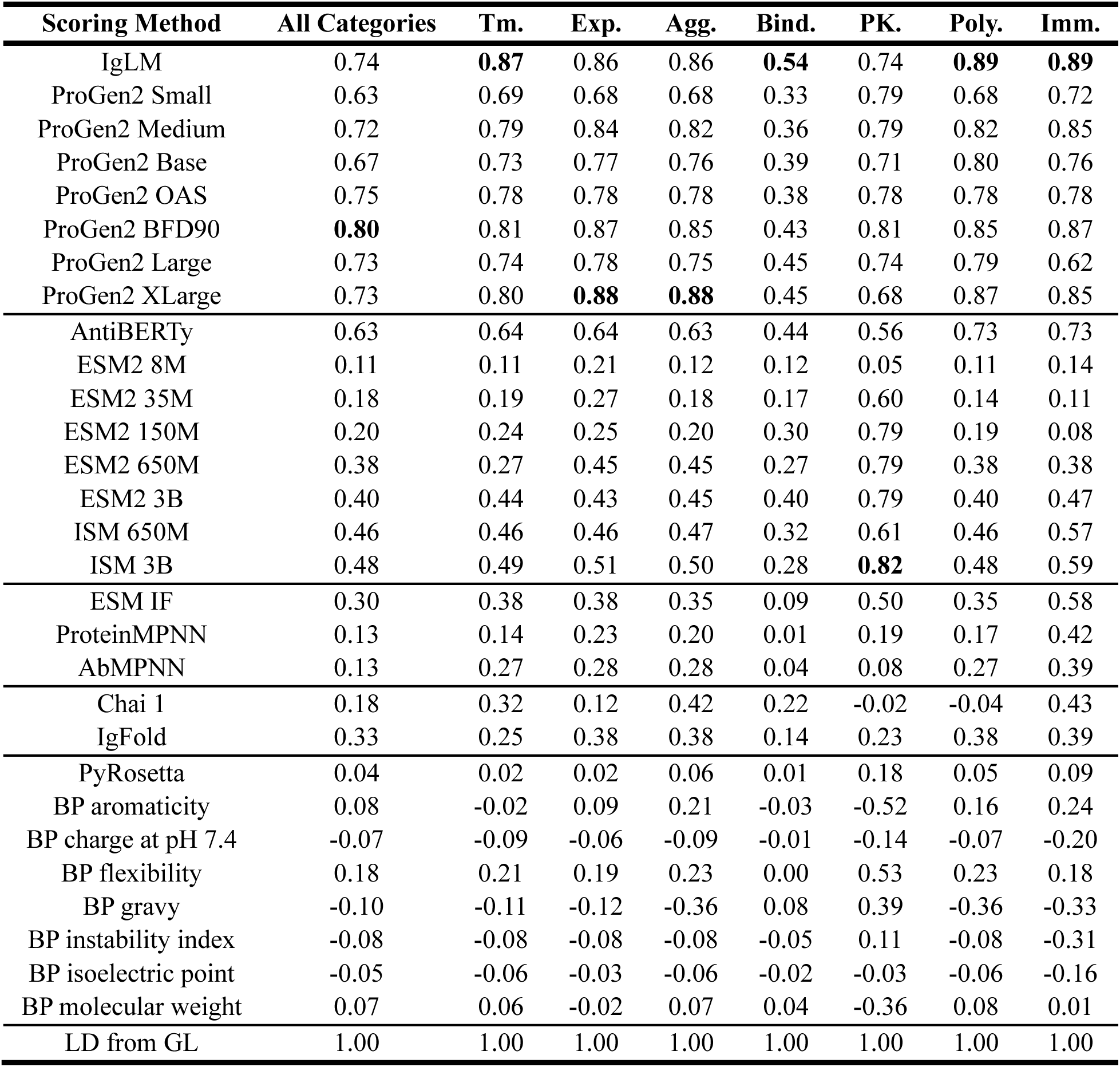
Zero-shot correlation with germline signal. Tabulated correlation between model confidence and distance from germline. A correlation approaching 1.0 indicates the model is highly confident in germline sequences and highly unconfident in sequences far from germline.

**Table S10:** Zero-shot correlation to germline signal across sequence-only and structure-informed models. Masked language modes without structure are AntiBERTY and the ESM2 suite. Masked language models with structure include the ISM suite. Inverse folding models are ESM-IF, ProteinMPNN, and AbMPNN. Structure predictors are Chai-1 and IgFold. Sequence-only masked language and causal language models are the ESM2 suite, AntiBERTY, IgLM, and ProGen2.

| Learned rep. | All Categories | Tm. | Exp. | Agg. | Bind. | PK. | Poly. | Imm. |
| --- | --- | --- | --- | --- | --- | --- | --- | --- |
| MLMs w/o structure | 0.32 | 0.31 | 0.37 | 0.34 | 0.28 | 0.60 | 0.32 | 0.32 |
| MLM w/ structure | 0.47 | 0.47 | 0.49 | 0.49 | 0.30 | **0.72** | 0.47 | 0.58 |
| Inverse folding | 0.19 | 0.26 | 0.30 | 0.28 | 0.05 | 0.26 | 0.26 | 0.46 |
| Structure predictor | 0.26 | 0.29 | 0.25 | 0.40 | 0.18 | 0.11 | 0.17 | 0.41 |
| MLMs and CLMs | **0.54** | **0.56** | **0.60** | **0.59** | **0.35** | 0.69 | **0.59** | **0.59** |

**Figure S4:**
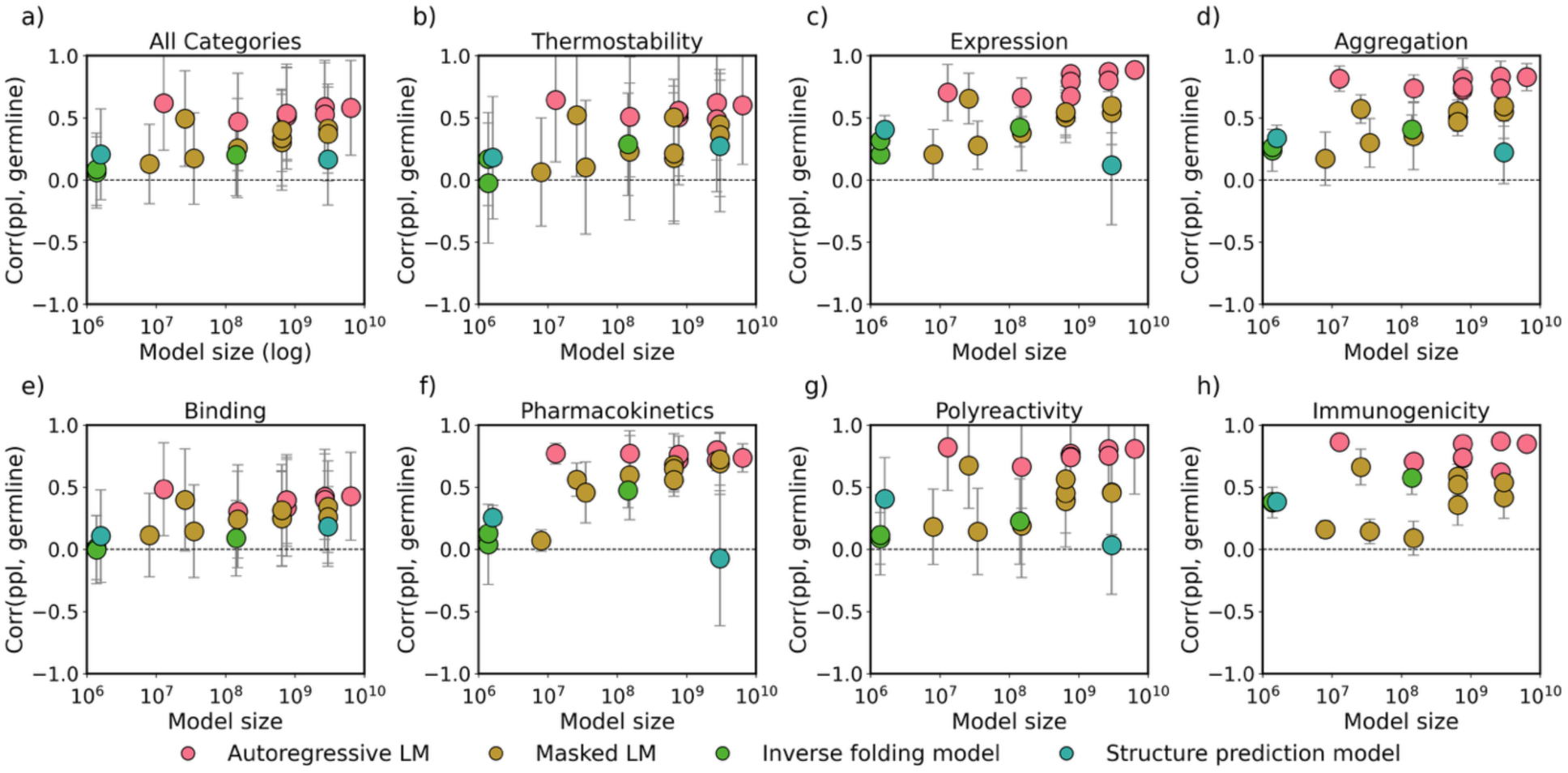
Zero-shot prediction correlations with germline compared to parameter size. Models are colored based on their architecture, and the *y*-axis displays the range of Spearman’s correlations between model confidence and the distance from germline. A correlation approaching 1.0 indicates the model is highly confident in germline sequences and highly unconfident in sequences far from germline. We only report Spearman’s correlations with datasets that exhibit at least five significant dataset-model p-values to ensure our conclusions are statistically sound.

**Figure S5:**
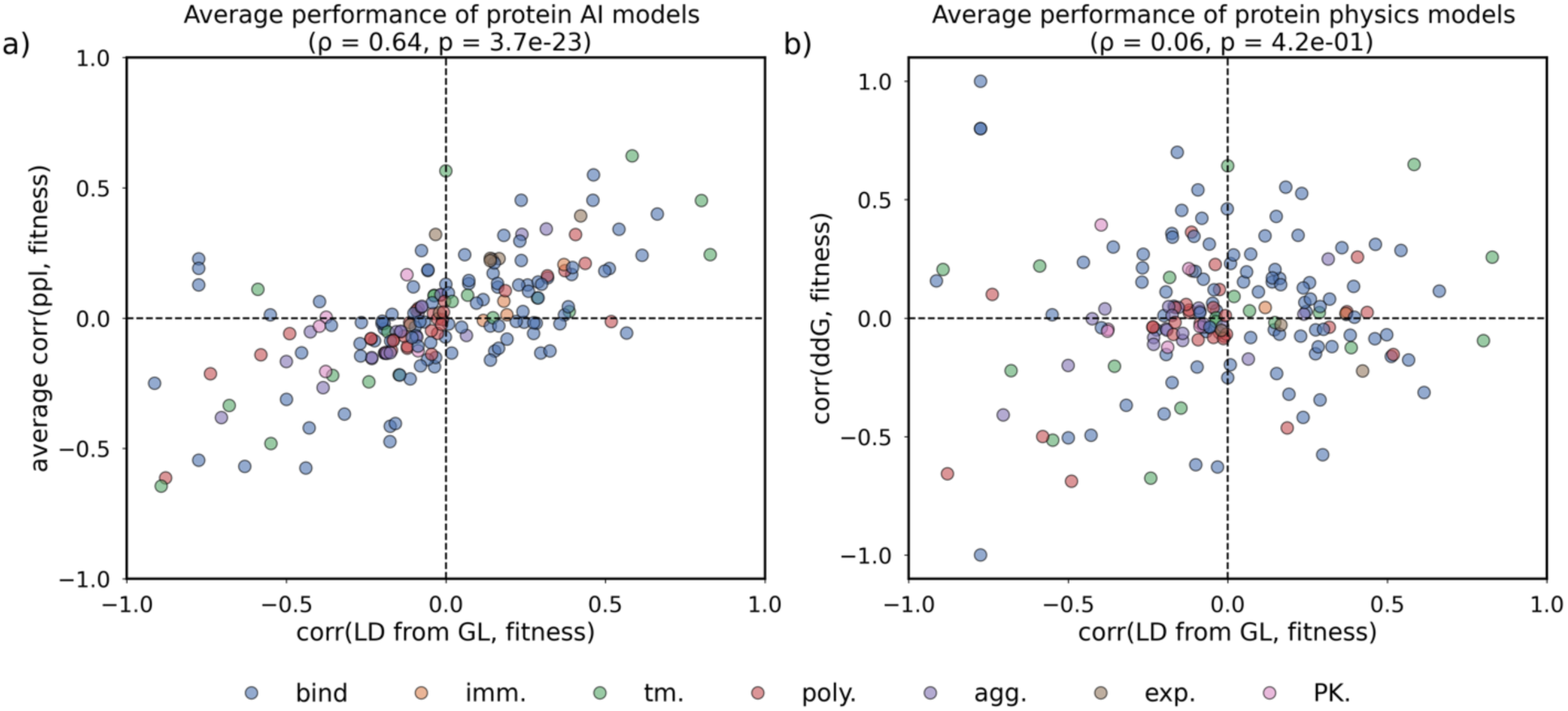
Average zero-shot prediction performance across datasets with varying directions of germline and fitness. Each point represents one developability dataset. The *x*-axis is the correlation between the distance from germline and the fitness, where a correlation approaching -1.0 indicates the sequences closer to germline have lower fitness, and a correlation approaching 1.0 indicates the sequences closer to germline have higher fitness. The *y*-axis is the average correlation of perplexity to fitness across all benchmarked protein AI models. a) Protein AI models, and b) PyRosetta. We only report Spearman’s correlations with five significant dataset-model p-values to ensure our conclusions are statistically sound.

#### Partial correlation calculation

The partial correlation coefficient^60^ measures the relationship between two variables (*A* , *B*) while eliminating influence of a third variable *C* .

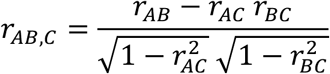

We calculate the partial correlation between the protein AI model perplexity (*A*) and the developability label (*B*) accounting for germline signal (*C* ). For a given developability dataset, we calculate the Spearman’s correlations between the AI model perplexity and the developability label (*r*_56_), between the AI model and germline signal (*r*_58_), and between the developability label and germline signal (*r*_68_).

**Table S11:**
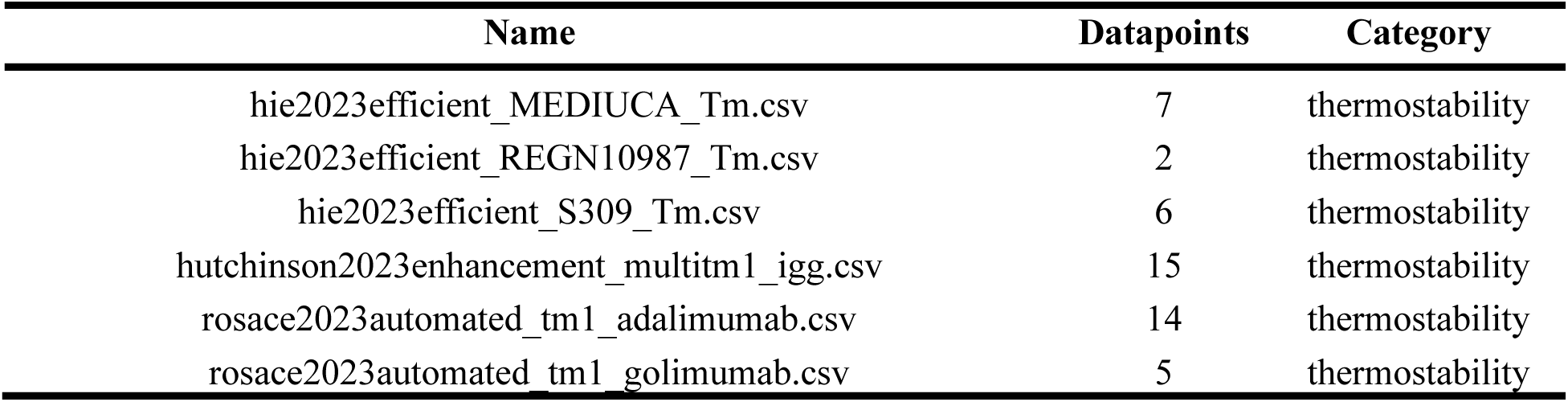
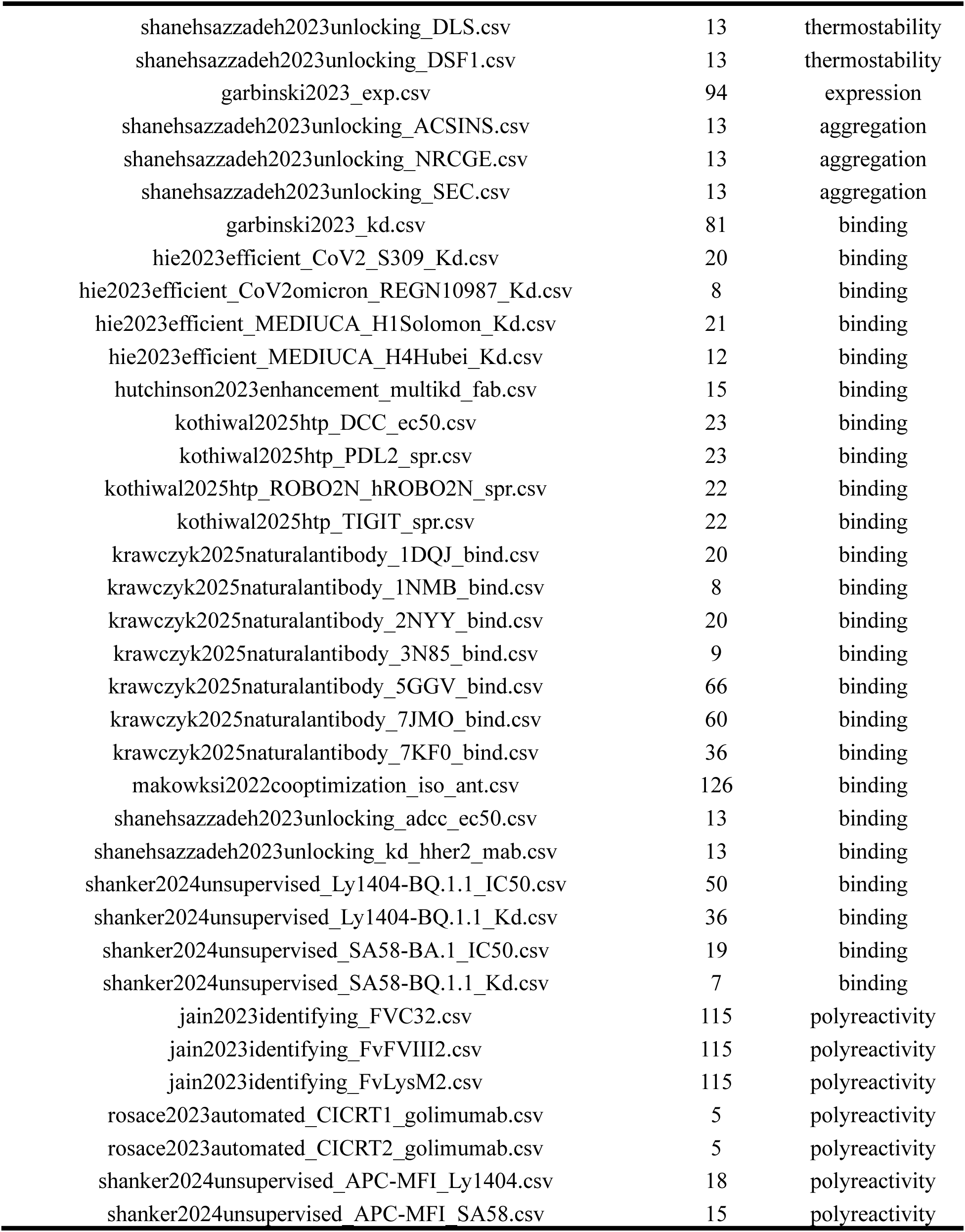
Partial correlation datasets. Datasets for which at least five models show statistically significant correlations and for which the mean Spearman correlation across models is at least 0.3.

**Table S12:** Partial correlations of protein AI models after accounting for germline signal.

| Scoring Method | Original corr. | Adjusted corr. | Diff. |
| --- | --- | --- | --- |
| IgLM | 0.50 | 0.26 | 0.47 |
| ProGen2 Small | 0.47 | 0.29 | 0.38 |
| ProGen2 Medium | 0.46 | 0.23 | 0.51 |
| ProGen2 Base | 0.48 | 0.30 | 0.37 |
| ProGen2 OAS | 0.48 | 0.23 | 0.52 |
| ProGen2 BFD90 | 0.48 | 0.23 | 0.52 |
| ProGen2 Large | 0.49 | 0.21 | 0.57 |
| ProGen2 XLarge | 0.47 | 0.17 | **0.63** |
| AntiBERTy | 0.49 | 0.30 | 0.39 |
| ESM2 8M | 0.46 | 0.39 | 0.16 |
| ESM2 35M | 0.53 | 0.38 | 0.27 |
| ESM2 150M | **0.54** | 0.43 | 0.21 |
| ESM2 650M | 0.50 | 0.35 | 0.31 |
| ESM2 3B | 0.49 | 0.32 | 0.35 |
| ISM 650M | 0.50 | 0.35 | 0.30 |
| ISM 3B | 0.52 | 0.37 | 0.28 |
| ESM IF | 0.46 | 0.38 | 0.18 |
| ProteinMPNN | 0.51 | 0.46 | 0.09 |
| AbMPNN | 0.51 | 0.41 | 0.20 |
| Chai 1 | 0.53 | 0.42 | 0.20 |
| IgFold | 0.52 | 0.36 | 0.31 |
| PyRosetta | 0.48 | 0.40 | 0.16 |
| BP aromaticity | 0.48 | 0.40 | 0.16 |
| BP charge at pH 7.4 | 0.48 | 0.45 | 0.06 |
| BP flexibility | 0.49 | 0.37 | 0.25 |
| BP instability index | 0.49 | 0.45 | 0.09 |
| BP isoelectric point | 0.51 | **0.48** | 0.05 |
| BP molecular weight | 0.46 | 0.38 | 0.17 |

### 10.5 Few-shot results

**Table S13:**
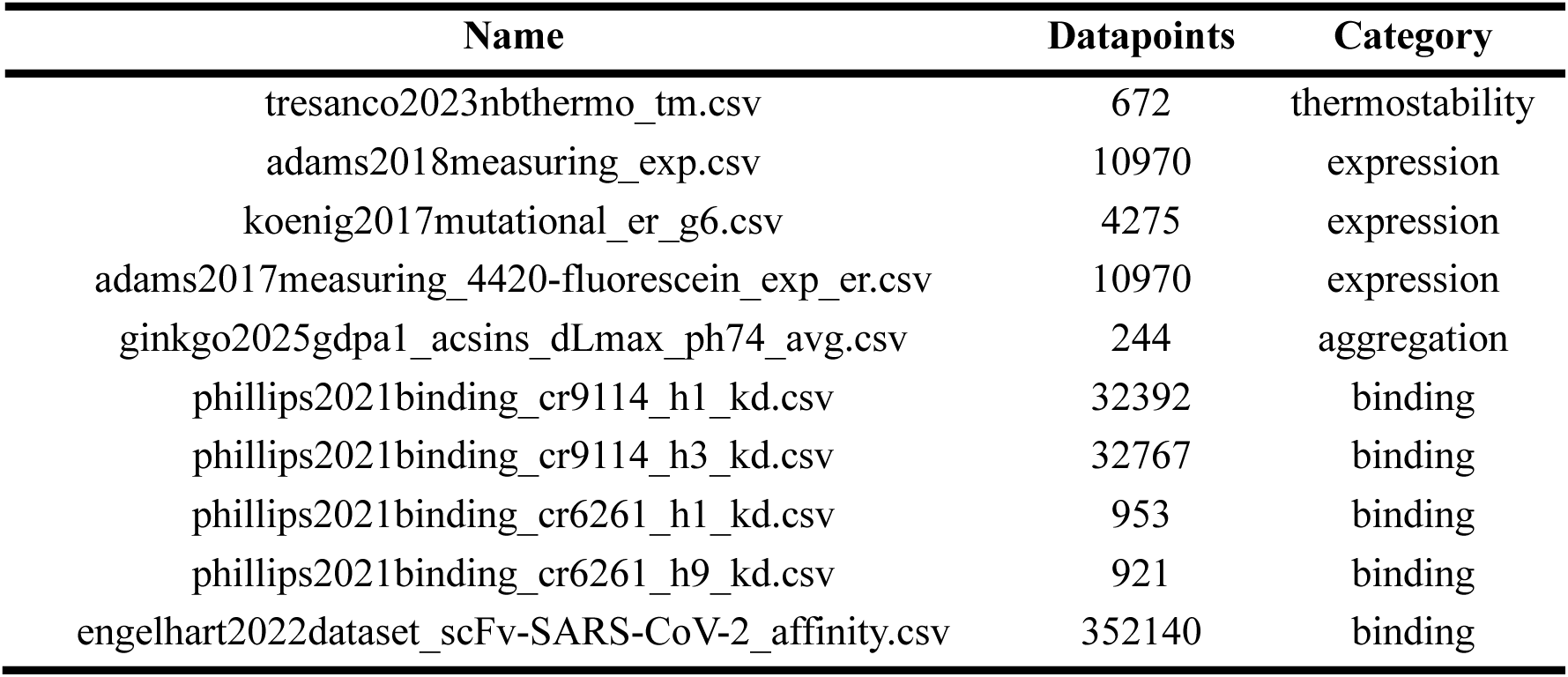
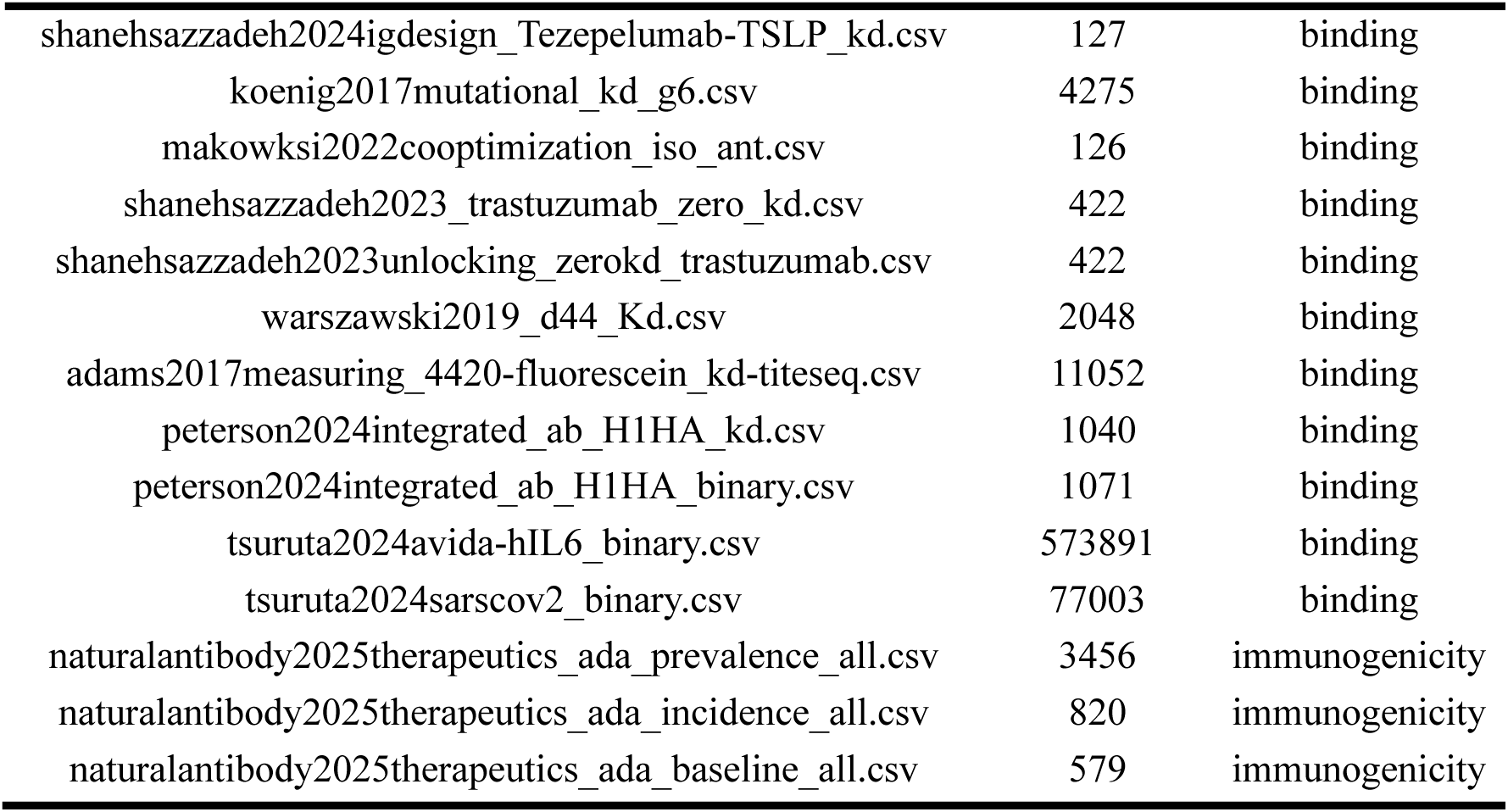
Datasets that pass the 5-significant-model cutoff. Datasets that satisfy the 5-significant-model cutoff and are used for the few-shot model analyses.

**Figure S6:**
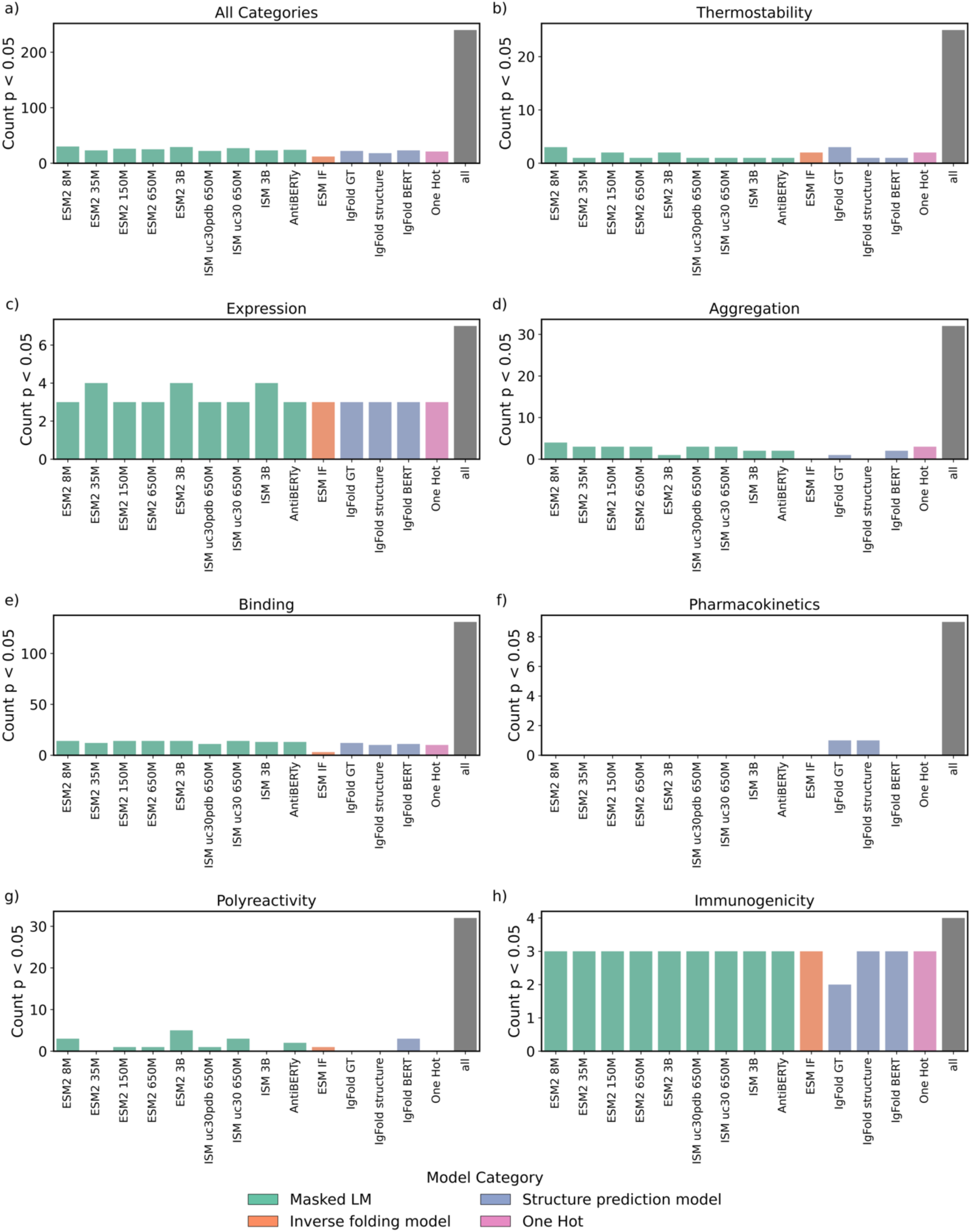
Count of significant correlations per few-shot model. Count of significant (p-value < 0.05) correlations each protein AI model had within each developability category. We only report Spearman’s correlations with five significant dataset-model p values to ensure our conclusions are statistically sound.

**Table S14:** Few-shot performance summary across all models. Tabulated correlation between model confidence and actual fitness score, averaged across all models. Some of the assay labels are inverted so that a positive correlation between fitness and assay value always indicates improved model performance.

| All Categories | Tm. | Exp. | Agg. | Bind. | Imm. |
| --- | --- | --- | --- | --- | --- |
| 0.49 | **0.61** | 0.53 | 0.32 | 0.43 | 0.44 |

**Table S15:** Few-shot performance summary for each individual model. Tabulated correlation between model confidence and actual fitness score, for each model.

| Scoring Method | All Categories | Tm. | Exp. | Agg. | Bind. | Imm. |
| --- | --- | --- | --- | --- | --- | --- |
| AntiBERTy | 0.54 | 0.60 | 0.55 | 0.52 | 0.47 | 0.45 |
| ESM2 8M | 0.51 | 0.57 | 0.55 | 0.25 | 0.39 | 0.47 |
| ESM2 35M | 0.50 | 0.70 | **0.56** | 0.28 | 0.42 | 0.47 |
| ESM2 150M | 0.55 | 0.61 | 0.55 | 0.48 | 0.52 | **0.49** |
| ESM2 650M | 0.51 | 0.63 | 0.54 | 0.40 | 0.48 | 0.43 |
| ESM2 3B | **0.56** | 0.63 | 0.54 | 0.27 | 0.56 | 0.44 |
| ISM uc30 650M | 0.55 | **0.75** | 0.55 | 0.42 | 0.55 | 0.44 |
| ISM uc30pdb 650M | 0.54 | 0.72 | 0.55 | **0.66** | 0.45 | 0.45 |
| ISM 3B | **0.56** | 0.74 | **0.56** | 0.37 | **0.61** | 0.45 |
| ESM IF | 0.29 | 0.50 | 0.46 | 0.18 | 0.20 | 0.47 |
| IgFold BERT | 0.48 | 0.54 | 0.55 | 0.43 | 0.41 | 0.47 |
| IgFold GT | 0.40 | 0.51 | 0.47 | -0.03 | 0.33 | 0.40 |
| IgFold structure | 0.41 | 0.51 | 0.54 | -0.03 | 0.37 | 0.34 |
| One Hot | 0.42 | 0.51 | 0.47 | 0.32 | 0.30 | 0.36 |

**Table S16:** Few-shot performance across architectures. Masked language models include AntiBERTy, the ESM2 suite, and the ISM suite. Inverse folding models include ESM IF. Structure predictors include IgFold.

| Architecture | All Categories | Tm. | Exp. | Agg. | Bind. | Imm. |
| --- | --- | --- | --- | --- | --- | --- |
| Masked LMs | **0.54** | **0.66** | **0.55** | **0.41** | **0.49** | 0.45 |
| Inverse folding | 0.29 | 0.50 | 0.46 | 0.18 | 0.20 | **0.47** |
| Structure predictors | 0.43 | 0.52 | 0.52 | 0.12 | 0.37 | 0.40 |
| One Hot | 0.42 | 0.51 | 0.47 | 0.32 | 0.30 | 0.36 |

**Table S17:** Few-shot performance across sequence-only and structure-informed models. Masked language modes without structure are AntiBERTY and the ESM2 suite. Masked language models with structure include the ISM suite. Inverse folding models are ESM-IF, ProteinMPNN, and AbMPNN. Structure predictors are Chai-1 and IgFold. Sequence-only masked language and causal language models are the ESM2 suite, AntiBERTY, IgLM, and ProGen2.

| Architecture | All Categories | Tm. | Exp. | Agg. | Bind. | Imm. |
| --- | --- | --- | --- | --- | --- | --- |
| MLM w/o structure | 0.53 | 0.62 | 0.55 | 0.37 | 0.47 | 0.46 |
| MLM w/ structure | **0.55** | **0.74** | **0.55** | **0.48** | **0.54** | 0.45 |
| Inverse folding | 0.29 | 0.50 | 0.46 | 0.18 | 0.20 | **0.47** |
| Structure predictors | 0.43 | 0.52 | 0.52 | 0.12 | 0.37 | 0.40 |
| One Hot | 0.42 | 0.51 | 0.47 | 0.32 | 0.30 | 0.36 |

**Table S18:** Few-shot performance across general protein and antibody-specific AI models. General protein AI models include the ESM2 suite, the ISM suite, and ESM IF. Antibody specific models include AntiBERTy and IgFold.

| Protein family | All Categories | Tm. | Exp. | Agg. | Bind. | Imm. |
| --- | --- | --- | --- | --- | --- | --- |
| General protein | **0.51** | **0.65** | **0.54** | **0.37** | **0.46** | **0.46** |
| Antibody specific | 0.46 | 0.54 | 0.53 | 0.22 | 0.39 | 0.42 |

**Figure S7:**
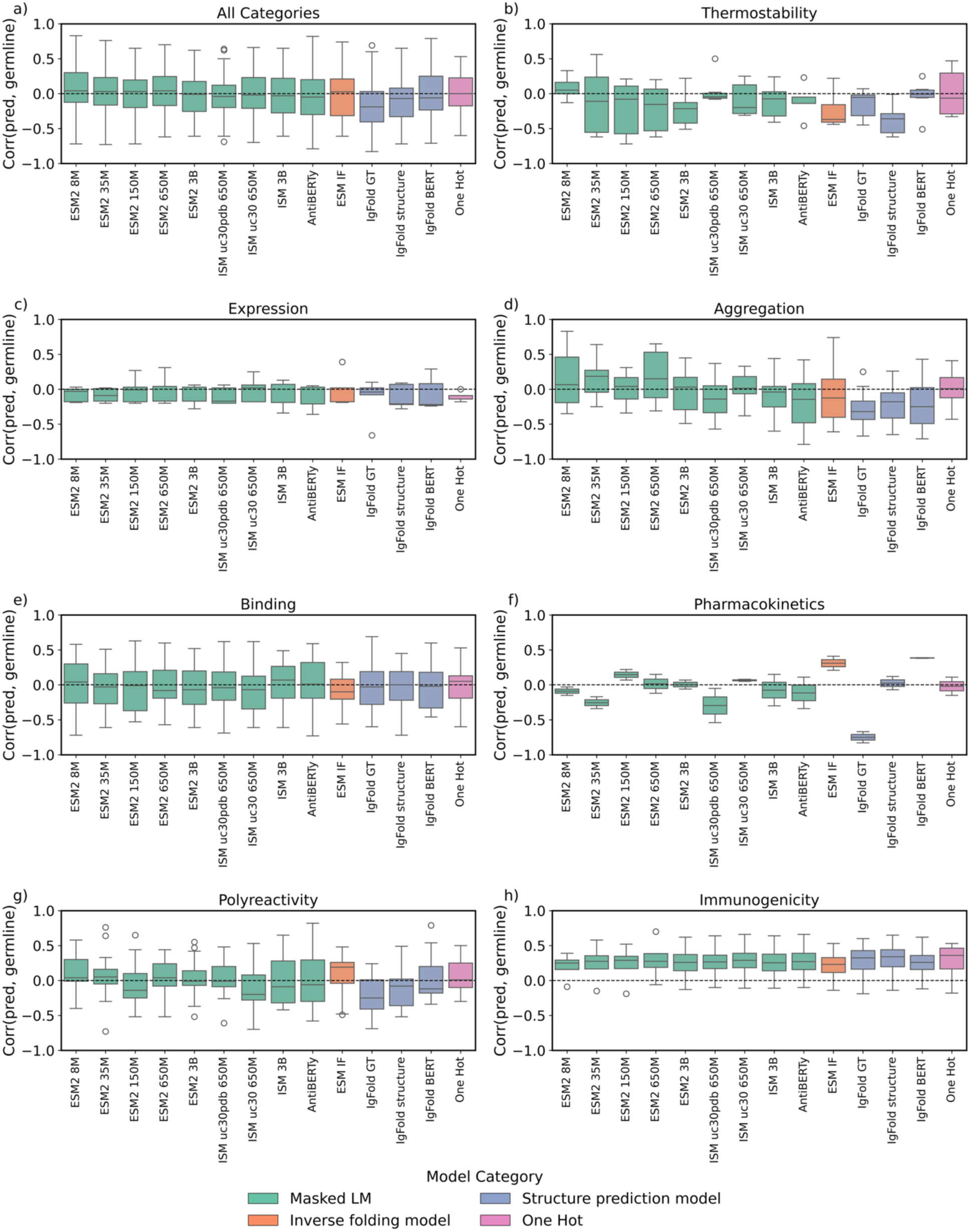
Distribution of few-shot prediction correlations with germline. Models are colored based on their architecture, and the *y*-axis displays the range of Spearman’s correlations between predicted developability and the distance from germline. A correlation approaching 1.0 indicates the model is highly confident in germline sequences and highly unconfident in sequences far from germline. We only report Spearman’s correlations with datasets that exhibit at least five significant dataset-model p-values to ensure our conclusions are statistically sound.

**Table S19:** Few-shot correlation with germline signal. Tabulated correlation between model confidence and distance from germline. A correlation approaching 1.0 indicates the model is highly confident in germline sequences and highly unconfident in sequences far from germline.

| Scoring Method | All Categories | Tm. | Exp. | Agg. | Bind. | PK. | Poly. | Imm. |
| --- | --- | --- | --- | --- | --- | --- | --- | --- |
| AntiBERTy | -0.05 | -0.05 | -0.21 | -0.15 | 0.01 | -0.12 | -0.06 | 0.27 |
| ESM2 8M | 0.04 | **0.05** | -0.03 | 0.07 | 0.04 | -0.09 | 0.04 | 0.25 |
| ESM2 35M | 0.03 | -0.11 | -0.09 | **0.19** | -0.03 | -0.26 | 0.05 | 0.28 |
| ESM2 150M | 0.03 | -0.08 | -0.01 | 0.04 | -0.01 | 0.15 | -0.14 | 0.29 |
| ESM2 650M | **0.04** | -0.16 | -0.17 | 0.15 | -0.08 | 0.02 | 0.04 | 0.28 |
| ESM2 3B | -0.01 | -0.22 | -0.17 | 0.03 | -0.07 | 0.01 | -0.01 | 0.26 |
| ISM uc30 650M | -0.02 | -0.20 | **0.02** | 0.02 | -0.07 | 0.07 | -0.20 | 0.29 |
| ISM uc30pdb 650M | -0.04 | -0.05 | -0.17 | -0.14 | -0.04 | -0.30 | -0.01 | 0.27 |
| ISM 3B | -0.03 | -0.08 | -0.19 | -0.04 | **0.07** | -0.08 | -0.09 | 0.26 |
| ESM IF | 0.03 | -0.37 | -0.18 | -0.13 | -0.10 | 0.31 | **0.19** | 0.23 |
| IgFold BERT | -0.06 | -0.01 | -0.22 | -0.25 | -0.02 | **0.39** | -0.12 | 0.26 |
| IgFold GT | -0.19 | -0.06 | -0.04 | -0.32 | -0.03 | -0.75 | -0.25 | 0.33 |
| IgFold structure | -0.07 | -0.36 | -0.21 | -0.18 | 0.00 | 0.02 | -0.08 | 0.34 |
| One Hot | 0.00 | -0.07 | -0.09 | 0.01 | 0.05 | -0.02 | 0.01 | **0.36** |

**Figure S8:**
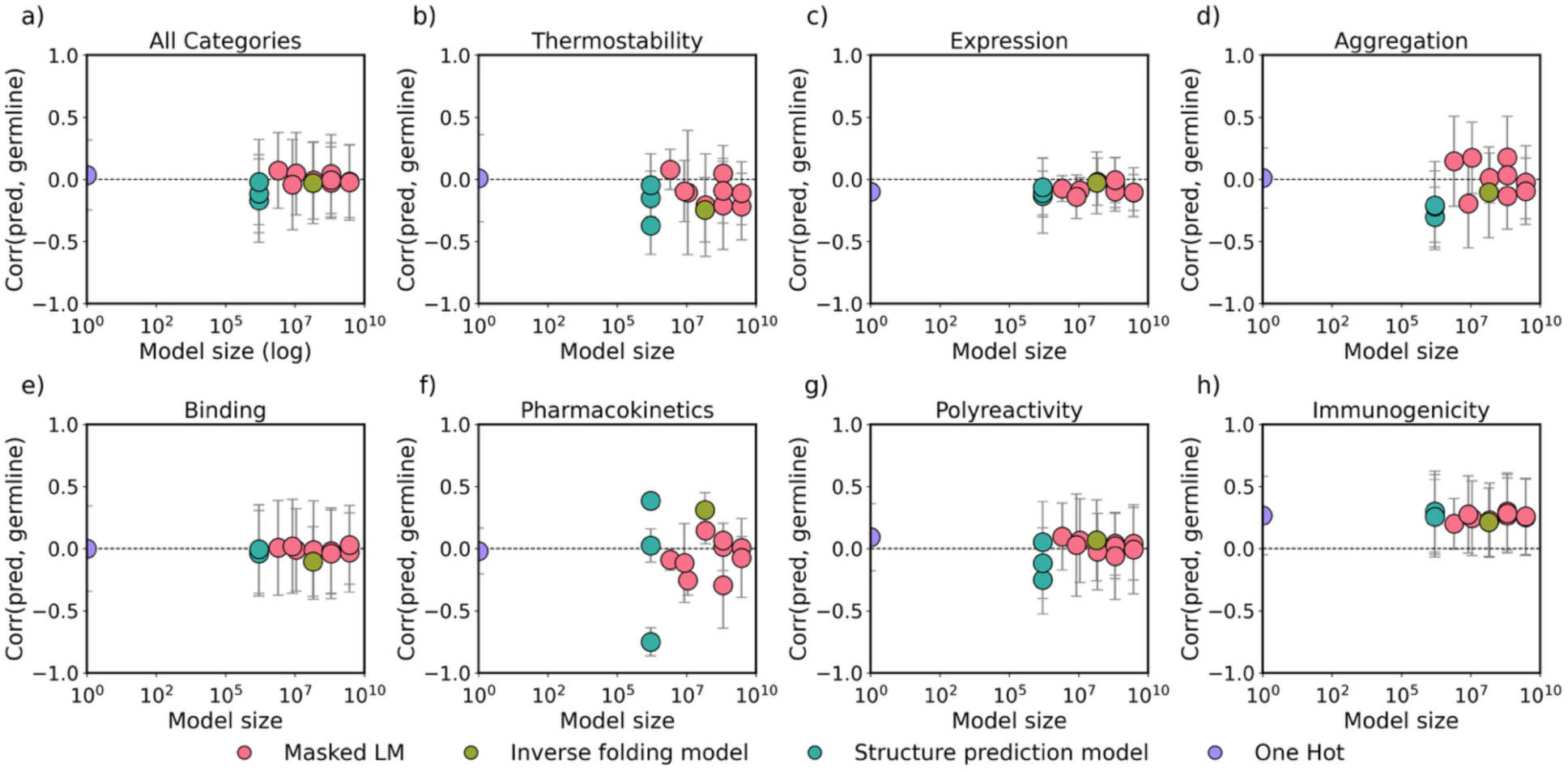
Distribution of few-shot prediction correlations with germline compared to parameter size. Models are colored based on their architecture, and the *y*-axis displays the range of Spearman’s correlations between predicted fitness and the distance from germline. A correlation approaching 1.0 indicates the model is highly confident in germline sequences and highly unconfident in sequences far from germline. We only report Spearman’s correlations with datasets that exhibit at least five significant dataset-model p-values to ensure our conclusions are statistically sound.

## Citations

1. Crescioli, S., Kaplon, H., Wang, L., Visweswaraiah, J., Kapoor, V. & Reichert, J. M. Antibodies to watch in 2025. MAbs 17, 2443538 (2025).

2. Bennett, N. R., Watson, J. L., Ragotte, R. J., Borst, A. J., See, D. L., Weidle, C., Biswas, R., Yu, Y., Shrock, E. L., Ault, R., Leung, P. J. Y., Huang, B., Goreshnik, I., Tam, J., Carr, K. D., Singer, B., Criswell, C., Wicky, B. I. M., Vafeados, D., et al. Atomically accurate de novo design of antibodies with RFdiffusion. bioRxivorg (2025) doi:10.1101/2024.03.14.585103.

3. Biswas, S. De novo design of epitope-specific antibodies against soluble and multipass membrane proteins with high specificity, developability, and function. Bioengineering (2025).

4. Boitreaud, J., Dent, J., Geisz, D., McPartlon, M., Meier, J., Qiao, Z., Rogozhnikov, A., Rollins, N., Wollenhaupt, P. & Wu, K. Zero-shot antibody design in a 24-well plate. Synthetic Biology (2025).

5. Frey, N. C., Hötzel, I., Stanton, S. D., Kelly, R., Alberstein, R. G., Makowski, E., Martinkus, K., Berenberg, D., Bevers, J., III, Bryson, T., Chan, P., Czubaty, A., D’Souza, T., Dwyer, H., Dziewulska, A., Fairman, J. W., Goodman, A., Hofmann, J., Isaacson, H., et al. Lab-in-the-loop therapeutic antibody design with deep learning. Bioengineering (2025).

6. Hie, B. L., Shanker, V. R., Xu, D., Bruun, T. U. J., Weidenbacher, P. A., Tang, S., Wu, W., Pak, J. E. & Kim, P. S. Efficient evolution of human antibodies from general protein language models. Nat. Biotechnol. 42, 275–283 (2024).

7. Shanker, V. R., Bruun, T. U. J., Hie, B. L. & Kim, P. S. Unsupervised evolution of protein and antibody complexes with a structure-informed language model. Science 385, 46–53 (2024).

8. Mazrooei, P., O’Neil, D., Izadi, S., Chen, B. & Ramanujan, S. Leveraging protein language and structural models for early prediction of antibodies with fast clearance. Pharmacology and Toxicology (2024).

9. Anapindi, K. D. B., Liu, K., Wang, W., Yu, Y., He, Y., Hsieh, E. J., Huang, Y. & Tomazela, D. Leveraging multi-modal feature learning for predictions of antibody viscosity. MAbs 17, 2490788 (2025).

10. Jain, T., Prinz, B., Marker, A., Michel, A., Reichel, K., Czepczor, V., Klieber, S., Sun, W., Kathuria, S., Oezguer Bruederle, S., Lange, C., Wahl, L., Starr, C., Masiero, A. & Avery, L. Assessment and incorporation of in vitro correlates to pharmacokinetic outcomes in antibody developability workflows. MAbs 16, 2384104 (2024).

11. Yu, X., Vangjeli, K., Prakash, A., Chhaya, M., Stanley, S. J., Cohen, N. & Huang, L. Protein language models enable prediction of polyreactivity of monospecific, bispecific, and heavy-chain-only antibodies. Antib. Ther. 7, 199–208 (2024).

12. Arsiwala, A., Bhatt, R., Yang, Y., Quintero Cadena, P., Anderson, K. C., Ao, X., van Niekerk, L., Rosenbaum, A., Bhatt, A., Smith, A., Grippo, L., Cao, X., Cohen, R., Patel, J., Allen, O., Faraj, A., Nandy, A., Hocking, J., Tural, B., et al. A high-throughput platform for biophysical antibody developability assessment to enable AI/ML model training. Biophysics (2025).

13. Chungyoun, M. & Gray, J. J. AI models for protein design are driving antibody engineering. Curr. Opin. Biomed. Eng. 28, 100473 (2023).

14. Joubbi, S., Micheli, A., Milazzo, P., Maccari, G., Ciano, G., Cardamone, D. & Medini, D. Antibody design using deep learning: from sequence and structure design to affinity maturation. Brief. Bioinform. 25, bbae307 (2024).

15. Jin, W., Wohlwend, J., Barzilay, R. & Jaakkola, T. Iterative refinement graph neural network for antibody sequence-structure co-design. arXiv [q-bio.BM] (2021).

16. Luo, S., Su, Y., Peng, X., Wang, S., Peng, J. & Ma, J. Antigen-specific antibody design and optimization with diffusion-based generative models for protein structures. bioRxiv (2022) doi:10.1101/2022.07.10.499510.

17. Kong, X., Huang, W. & Liu, Y. End-to-end full-atom antibody design. arXiv [q-bio.BM] (2023).

18. Xu, B., Wang, Y., Chen, W. & Shan, S. AntibodyFlow: Normalizing flow model for designing antibody complementarity-determining regions. arXiv [cs.LG] (2024).

19. Wang, R., Wu, F., Gao, X., Wu, J., Zhao, P. & Yao, J. IgGM: A Generative Model for Functional Antibody and Nanobody Design. Bioinformatics (2024).

20. Ren, M., He, Z. & Zhang, H. Multi-objective antibody design with constrained preference optimization. (2024).

21. Chen, C., Herpoldt, K.-L., Zhao, C., Wang, Z., Collins, M., Shang, S. & Benson, R. AffinityFlow: Guided flows for antibody affinity maturation. arXiv [cs.LG] (2025).

22. Olsen, T. H., Boyles, F. & Deane, C. M. Observed Antibody Space: A diverse database of cleaned, annotated, and translated unpaired and paired antibody sequences. Protein Sci. 31, 141–146 (2022).

23. Dudzic, P., Chomicz, D., Bielska, W., Jaszczyszyn, I., Zieliński, M., Janusz, B., Wróbel, S., Pannérer, M.-M. L., Philips, A., Ponraj, P., Kumar, S. & Krawczyk, K. Conserved heavy/light contacts and germline preferences revealed by a large-scale analysis of natively paired human antibody sequences and structural data. Immunology (2024).

24. Schneider, C., Raybould, M. I. J. & Deane, C. M. SAbDab in the age of biotherapeutics: updates including SAbDab-nano, the nanobody structure tracker. Nucleic Acids Res. 50, D1368–D1372 (2022).

25. Zhou, N., Jiang, Y., Bergquist, T. R., Lee, A. J., Kacsoh, B. Z., Crocker, A. W., Lewis, K. A., Georghiou, G., Nguyen, H. N., Hamid, M. N., Davis, L., Dogan, T., Atalay, V., Rifaioglu, A. S., Dalkıran, A., Cetin Atalay, R., Zhang, C., Hurto, R. L., Freddolino, P. L., et al. The CAFA challenge reports improved protein function prediction and new functional annotations for hundreds of genes through experimental screens. Genome Biol. 20, 244 (2019).

26. Rao, R., Bhattacharya, N., Thomas, N., Duan, Y., Chen, X., Canny, J., Abbeel, P. & Song, Y. S. Evaluating protein transfer learning with TAPE. Adv. Neural Inf. Process. Syst. 32, 9689–9701 (2019).

27. Dallago, C., Mou, J., Johnston, K. E., Wittmann, B. J., Bhattacharya, N., Goldman, S., Madani, A. & Yang, K. K. FLIP: Benchmark tasks in fitness landscape inference for proteins. bioRxiv (2021) doi:10.1101/2021.11.09.467890.

28. Notin, P., Kollasch, A. W., Ritter, D., van Niekerk, L., Paul, S., Spinner, H., Rollins, N., Shaw, A., Weitzman, R., Frazer, J., Dias, M., Franceschi, D., Orenbuch, R., Gal, Y. & Marks, D. S. ProteinGym: Large-scale benchmarks for protein design and fitness prediction. bioRxivorg (2023) doi:10.1101/2023.12.07.570727.

29. Garcia, M., Dixit, S. M. & Rocklin, G. J. Evaluating zero-shot prediction of protein design success by AlphaFold, ESMFold, and ProteinMPNN. Bioinformatics (2025).

30. Ma, Z. & Bethel, N. P. RemoteFoldSet: Benchmarking Structural Awareness of Protein Language Models. Biochemistry (2025).

31. Shanehsazzadeh, A., McPartlon, M., Kasun, G., Steiger, A. K., Sutton, J. M., Yassine, E., McCloskey, C., Haile, R., Shuai, R., Alverio, J., Rakocevic, G., Levine, S., Cejovic, J., Gutierrez, J. M., Morehead, A., Dubrovskyi, O., Chung, C., Luton, B. K., Diaz, N., et al. Unlocking de novo antibody design with generative artificial intelligence. Synthetic Biology (2023).

32. Porebski, B. T., Balmforth, M., Browne, G., Riley, A., Jamali, K., Fürst, M. J. L. J., Velic, M., Buchanan, A., Minter, R., Vaughan, T. & Holliger, P. Rapid discovery of high-affinity antibodies via massively parallel sequencing, ribosome display and affinity screening. *Nat*. Biomed. Eng. 8, 214–232 (2024).

33. Uçar, T., Malherbe, C. & Gonzalez, F. Exploring Log-Likelihood Scores for Ranking Antibody Sequence Designs. Bioinformatics (2024).

34. Liu, C., Pelissier, A., Shao, Y., Denzler, L., Martin, A. C. R., Paige, B. & Martinez, M. R. AbRank: A benchmark dataset and metric-learning framework for antibody-antigen affinity ranking. arXiv [q-bio.BM] (2025).

35. Janusz, B., Chomicz, D., Demharter, S., Arts, M., de Kanter, J., Wilke, Y., Britze, H., Wrobel, S., Gawłowski, T., Dudzic, P., Ukkivi, K., Peil, L., Spreafico, R. & Krawczyk, K. AbDesign: Database of point mutants of antibodies with associated structures reveals poor generalization of binding predictions from machine learning models. Bioinformatics (2025).

36. Cia, G., Li, D., Poblete, S., Rooman, M. & Pucci, F. AbAgym: a well-curated dataset for the mutational analysis of antibody-antigen complexes. bioRxiv 2025.07.15.664862 (2025).

37. Zhao, X., Tang, Y.-C., Singh, A., Cantu, V. J., An, K., Lee, J., Stogsdill, A. E., Hamdi, I. M., Ramesh, A. K., An, Z., Jiang, X. & Kim, Y. AbBiBench: A benchmark for antibody binding affinity maturation and design. arXiv [q-bio.QM] (2025) doi:10.48550/arXiv.2506.04235.

38. Chungyoun, M., Ruffolo, J. & Gray, J. FLAb: Benchmarking deep learning methods for antibody fitness prediction. bioRxiv 2024.01.13.575504 (2024) doi:10.1101/2024.01.13.575504.

39. Jain, T., Sun, T., Durand, S., Hall, A., Houston, N. R., Nett, J. H., Sharkey, B., Bobrowicz, B., Caffry, I., Yu, Y., Cao, Y., Lynaugh, H., Brown, M., Baruah, H., Gray, L. T., Krauland, E. M., Xu, Y., Vásquez, M. & Wittrup, K. D. Biophysical properties of the clinical-stage antibody landscape. Proc. Natl. Acad. Sci. U. S. A. 114, 944–949 (2017).

40. Shuai, R. W., Ruffolo, J. A. & Gray, J. J. IgLM: Infilling language modeling for antibody sequence design. Cell Syst. 14, 979–989.e4 (2023).

41. Nijkamp, E., Ruffolo, J. A., Weinstein, E. N., Naik, N. & Madani, A. ProGen2: Exploring the boundaries of protein language models. Cell Syst. 14, 968–978.e3 (2023).

42. Ruffolo, J. A., Gray, J. J. & Sulam, J. Deciphering antibody affinity maturation with language models and weakly supervised learning. arXiv [q-bio.BM] (2021).

43. Lin, Z., Akin, H., Rao, R., Hie, B., Zhu, Z., Lu, W., Smetanin, N., Verkuil, R., Kabeli, O., Shmueli, Y., Dos Santos Costa, A., Fazel-Zarandi, M., Sercu, T., Candido, S. & Rives, A. Evolutionary-scale prediction of atomic-level protein structure with a language model. Science 379, 1123–1130 (2023).

44. Ouyang-Zhang, J., Gong, C., Zhao, Y., Krähenbühl, P., Klivans, A. R. & Diaz, D. J. Distilling structural representations into protein sequence models. bioRxiv (2024) doi:10.1101/2024.11.08.622579.

45. Dreyer, F. A., Cutting, D., Schneider, C., Kenlay, H. & Deane, C. M. Inverse folding for antibody sequence design using deep learning. arXiv [q-bio.BM] (2023).

46. Hsu, C., Verkuil, R., Liu, J., Lin, Z., Hie, B., Sercu, T., Lerer, A. & Rives, A. Learning inverse folding from millions of predicted structures. bioRxiv (2022) doi:10.1101/2022.04.10.487779.

47. Dauparas, J., Anishchenko, I., Bennett, N., Bai, H., Ragotte, R. J., Milles, L. F., Wicky, B. I. M., Courbet, A., de Haas, R. J., Bethel, N., Leung, P. J. Y., Huddy, T. F., Pellock, S., Tischer, D., Chan, F., Koepnick, B., Nguyen, H., Kang, A., Sankaran, B., et al. Robust deep learning-based protein sequence design using ProteinMPNN. Science 378, 49–56 (2022).

48. Ruffolo, J. A., Chu, L.-S., Mahajan, S. P. & Gray, J. J. Fast, accurate antibody structure prediction from deep learning on massive set of natural antibodies. Nat. Commun. 14, 2389 (2023).

49. Discovery, C., Boitreaud, J., Dent, J., McPartlon, M., Meier, J., Reis, V., Rogozhnikov, A. & Wu, K. Chai-1: Decoding the molecular interactions of life. bioRxiv (2024) doi:10.1101/2024.10.10.615955.

50. Leaver-Fay, A., Tyka, M., Lewis, S. M., Lange, O. F., Thompson, J., Jacak, R., Kaufman, K., Renfrew, P. D., Smith, C. A., Sheffler, W., Davis, I. W., Cooper, S., Treuille, A., Mandell, D. J., Richter, F., Ban, Y.-E. A., Fleishman, S. J., Corn, J. E., Kim, D. E., et al. ROSETTA3: an object-oriented software suite for the simulation and design of macromolecules. Methods Enzymol. 487, 545–574 (2011).

51. Cock, P. J. A., Antao, T., Chang, J. T., Chapman, B. A., Cox, C. J., Dalke, A., Friedberg, I., Hamelryck, T., Kauff, F., Wilczynski, B. & de Hoon, M. J. L. Biopython: freely available Python tools for computational molecular biology and bioinformatics. Bioinformatics 25, 1422–1423 (2009).

52. Hutchinson, M., Ruffolo, J. A., Haskins, N., Iannotti, M., Vozza, G., Pham, T., Mehzabeen, N., Shandilya, H., Rickert, K., Croasdale-Wood, R., Damschroder, M., Fu, Y., Dippel, A., Gray, J. J. & Kaplan, G. Toward enhancement of antibody thermostability and affinity by computational design in the absence of antigen. MAbs 16, 2362775 (2024).

53. Stüber, J. C., Rechberger, K. F., Miladinović, S. M., Pöschinger, T., Zimmermann, T., Villenave, R., Eigenmann, M. J., Kraft, T. E., Shah, D. K., Kettenberger, H. & Richter, W. F. Impact of charge patches on tumor disposition and biodistribution of therapeutic antibodies. AAPS Open 8, (2022).

54. Ausserwöger, H., Krainer, G., Welsh, T. J., Thorsteinson, N., de Csilléry, E., Sneideris, T., Schneider, M. M., Egebjerg, T., Invernizzi, G., Herling, T. W., Lorenzen, N. & Knowles, T. P. J. Surface patches induce nonspecific binding and phase separation of antibodies. Proc. Natl. Acad. Sci. U. S. A. 120, e2210332120 (2023).

55. Kaplan, J., McCandlish, S., Henighan, T., Brown, T. B., Chess, B., Child, R., Gray, S., Radford, A., Wu, J. & Amodei, D. Scaling laws for neural language models. arXiv [cs.LG] (2020).

56. Brown, T. B., Mann, B., Ryder, N., Subbiah, M., Kaplan, J., Dhariwal, P., Neelakantan, A., Shyam, P., Sastry, G., Askell, A., Agarwal, S., Herbert-Voss, A., Krueger, G., Henighan, T., Child, R., Ramesh, A., Ziegler, D. M., Wu, J., Winter, C., et al. Language Models are Few-Shot Learners. arXiv [cs.CL] (2020).

57. Radford, A., Wu, J., Child, R., Luan, D., Amodei, D. & Sutskever, I. Language Models are Unsupervised Multitask Learners. (2019).

58. Hou, C., Liu, D., Zafar, A. & Shen, Y. Understanding Language Model Scaling on Protein Fitness Prediction. Bioinformatics (2025).

59. Olsen, T. H., Moal, I. H. & Deane, C. M. Addressing the antibody germline bias and its effect on language models for improved antibody design. Bioinformatics 40, btae618 (2024).

60. Wang, J. Partial Correlation Coefficient. in Encyclopedia of Systems Biology 1634–1635 (Springer New York, 2013).

61. Weinstein, E. N., Amin, A. N., Frazer, J. & Marks, D. S. Non-identifiability and the Blessings of Misspecification in Models of Molecular Fitness. Evolutionary Biology (2022).

62. Ding, F. & Steinhardt, J. Protein language models are biased by unequal sequence sampling across the tree of life. Bioinformatics (2024).

63. Ucar, T. & Sormanni, P. BLOSUM is all you learn - generative antibody models reflect evolutionary priors. bioRxiv (2025) doi:10.1101/2025.10.26.684652.

64. Hallee, L., Peleg, T., Rafailidis, N. & Gleghorn, J. P. Protein Language Models are Accidental Taxonomists. Bioinformatics (2025).

65. Harding, F. A., Stickler, M. M., Razo, J. & DuBridge, R. B. The immunogenicity of humanized and fully human antibodies: residual immunogenicity resides in the CDR regions. MAbs 2, 256–265 (2010).

66. Hu, Z., Cohen, S. & Swanson, S. J. The immunogenicity of human-origin therapeutic antibodies are associated with V gene usage. Front. Immunol. 14, 1237754 (2023).

67. Gutierrez, Y. M. & Rocklin, G. J. Structural and energetic analysis of stabilizing indel mutations. Biophysics (2024).

68. Bikias, T., Stamkopoulos, E. & Reddy, S. T. PLMFit: benchmarking transfer learning with protein language models for protein engineering. Brief. Bioinform. 26, bbaf381 (2025).

69. Lu, W., Luu, R. K. & Buehler, M. J. Fine-tuning large language models for domain adaptation: Exploration of training strategies, scaling, model merging and synergistic capabilities. arXiv [cs.CL] (2024).

70. Zhu, C., Xu, B., Wang, Q., Zhang, Y. & Mao, Z. On the calibration of large language models and alignment. arXiv [cs.CL] (2023).

71. Dieckhaus, H., Brocidiacono, M., Randolph, N. Z. & Kuhlman, B. Transfer learning to leverage larger datasets for improved prediction of protein stability changes. Proc. Natl. Acad. Sci. U. S. A. 121, e2314853121 (2024).

72. Mialland, A., Fukunaga, S., Katsuki, R., Dong, Y., Yamaguchi, H. & Saito, Y. Fitness translocation: improving variant effect prediction with biologically-grounded data augmentation. Bioinformatics (2024).

73. Minot, M. & Reddy, S. T. Meta learning addresses noisy and under-labeled data in machine learning-guided antibody engineering. Cell Syst. 15, 4–18.e4 (2024).

74. Almeida, D. S., Almeida, M. V., Sampaio, J. V., Gaieta, E. M., Costa, A. H. S., Rabelo, F. F. A., Cavalcante, C. L., Sartori, G. R. & Silva, J. H. M. AbSet: A standardized data set of antibody structures for machine learning applications. J. Chem. Inf. Model. 65, 4767–4774 (2025).

75. Kamat, V., Rafique, A., Huang, T., Olsen, O. & Olson, W. The impact of different human IgG capture molecules on the kinetics analysis of antibody-antigen interaction. Anal. Biochem. 593, 113580 (2020).

76. Hsu, C., Nisonoff, H., Fannjiang, C. & Listgarten, J. Learning protein fitness models from evolutionary and assay-labeled data. Nat. Biotechnol. 40, 1114–1122 (2022).

77. Hummer, A. M., Schneider, C., Chinery, L. & Deane, C. M. Investigating the volume and diversity of data needed for generalizable antibody-antigen ΔΔG prediction. Nat. Comput. Sci. 1–13 (2025).

78. Kirby, M. B., Petersen, B. M., Faris, J. G., Kells, S. P., Sprenger, K. G. & Whitehead, T. A. Retrospective SARS-CoV-2 human antibody development trajectories are largely sparse and permissive. Proc. Natl. Acad. Sci. U. S. A. 122, e2412787122 (2025).

79. Dudzic, P. & Krawczyk, K. Herding cats: predicting immunogenicity from heterogeneous clinical trials data. bioRxiv (2025) doi:10.1101/2025.10.14.682375.

80. Jain, T., Boland, T. & Vásquez, M. Identifying developability risks for clinical progression of antibodies using high-throughput in vitro and in silico approaches. MAbs 15, 2200540 (2023).

81. Jetha, A., Thorsteinson, N., Jmeian, Y., Jeganathan, A., Giblin, P. & Fransson, J. Homology modeling and structure-based design improve hydrophobic interaction chromatography behavior of integrin binding antibodies. MAbs 10, 890–900 (2018).

82. Kraft, T. E., Richter, W. F., Emrich, T., Knaupp, A., Schuster, M., Wolfert, A. & Kettenberger, H. Heparin chromatography as an in vitro predictor for antibody clearance rate through pinocytosis. MAbs 12, 1683432 (2020).

83. Phillips, A. M., Lawrence, K. R., Moulana, A., Dupic, T., Chang, J., Johnson, M. S., Cvijovic, I., Mora, T., Walczak, A. M. & Desai, M. M. Binding affinity landscapes constrain the evolution of broadly neutralizing anti-influenza antibodies. Elife 10, e71393 (2021).

84. Engelhart, E., Emerson, R., Shing, L., Lennartz, C., Guion, D., Kelley, M., Lin, C., Lopez, R., Younger, D. & Walsh, M. E. A dataset comprised of binding interactions for 104,972 antibodies against a SARS-CoV-2 peptide. Sci. Data 9, 653 (2022).

85. Shanehsazzadeh, A., Alverio, J., Kasun, G., Levine, S., Calman, I., Khan, J. A., Chung, C., Diaz, N., Luton, B. K., Tarter, Y., McCloskey, C., Bateman, K. B., Carter, H., Chapman, D., Consbruck, R., Jaeger, A., Kohnert, C., Kopec-Belliveau, G., Sutton, J. M., et al. IgDesign: In vitro validated antibody design against multiple therapeutic antigens using inverse folding. Synthetic Biology (2023).

86. Koenig, P., Lee, C. V., Walters, B. T., Janakiraman, V., Stinson, J., Patapoff, T. W. & Fuh, G. Mutational landscape of antibody variable domains reveals a switch modulating the interdomain conformational dynamics and antigen binding. Proc. Natl. Acad. Sci. U. S. A. 114, E486–E495 (2017).

87. Makowski, E. K., Kinnunen, P. C., Huang, J., Wu, L., Smith, M. D., Wang, T., Desai, A. A., Streu, C. N., Zhang, Y., Zupancic, J. M., Schardt, J. S., Linderman, J. J. & Tessier, P. M. Co-optimization of therapeutic antibody affinity and specificity using machine learning models that generalize to novel mutational space. Nat. Commun. 13, 3788 (2022).

88. Rawat, P., Sharma, D., Prabakaran, R., Ridha, F., Mohkhedkar, M., Janakiraman, V. & Gromiha, M. M. Ab-CoV: a curated database for binding affinity and neutralization profiles of coronavirus-related antibodies. Bioinformatics 38, 4051–4052 (2022).

89. Rosace, A., Bennett, A., Oeller, M., Mortensen, M. M., Sakhnini, L., Lorenzen, N., Poulsen, C. & Sormanni, P. Automated optimisation of solubility and conformational stability of antibodies and proteins. Nat. Commun. 14, 1937 (2023).

90. Warszawski, S., Borenstein Katz, A., Lipsh, R., Khmelnitsky, L., Ben Nissan, G., Javitt, G., Dym, O., Unger, T., Knop, O., Albeck, S., Diskin, R., Fass, D., Sharon, M. & Fleishman, S. J. Optimizing antibody affinity and stability by the automated design of the variable light-heavy chain interfaces. PLoS Comput. Biol. 15, e1007207 (2019).

91. Zimmermann, J., Oakman, E. L., Thorpe, I. F., Shi, X., Abbyad, P., Brooks, C. L., 3rd, Boxer, S. G. & Romesberg, F. E. Antibody evolution constrains conformational heterogeneity by tailoring protein dynamics. Proc. Natl. Acad. Sci. U. S. A. 103, 13722–13727 (2006).

92. Adams, R. M., Mora, T., Walczak, A. M. & Kinney, J. B. Measuring the sequence-affinity landscape of antibodies with massively parallel titration curves. Elife 5, e23156 (2016).

93. Petersen, B. M., Kirby, M. B., Chrispens, K. M., Irvin, O. M., Strawn, I. K., Haas, C. M., Walker, A. M., Baumer, Z. T., Ulmer, S. A., Ayala, E., Rhodes, E. R., Guthmiller, J. J., Steiner, P. J. & Whitehead, T. A. An integrated technology for quantitative wide mutational scanning of human antibody Fab libraries. Nat. Commun. 15, 3974 (2024).

94. Erasmus, M. F., Spector, L., Ferrara, F., DiNiro, R., Pohl, T. J., Perea-Schmittle, K., Wang, W., Tessier, P. M., Richardson, C., Turner, L., Kumar, S., Bedinger, D., Sormanni, P., Fernández-Quintero, M. L., Ward, A. B., Loeffler, J. R., Swanson, O. M., Deane, C. M., Raybould, M. I. J., et al. AIntibody: an experimentally validated in silico antibody discovery design challenge. Nat. Biotechnol. 42, 1637–1642 (2024).

95. Kothiwal, D., Kollasch, A. W., Hollmer, N., Ghosh, A., Zhang, R., Anuganti, M., Paul, S. B., Zagar, Y., Abdollahi, M., Anderson, Z., Belay, F., Salotto, M., Ulmer, S., Abdelalim, Y. A., Kumar, S., Vangala, M., Yang, C., Chedotal, A., Jardine, J. G., et al. High-Throughput Machine Learning-Aided Antibody Discovery for Cell Surface Antigens. Biophysics (2025).

96. Marks, C., Hummer, A. M., Chin, M. & Deane, C. M. Humanization of antibodies using a machine learning approach on large-scale repertoire data. Bioinformatics 37, 4041–4047 (2021).

97. [No title]. https://naturalantibody.com/therapeutic-antibody-database/.

98. Sulea, T. Humanization of camelid single-domain antibodies. Methods Mol. Biol. 2446, 299–312 (2022).

99. Valdés-Tresanco, M. S., Valdés-Tresanco, M. E., Molina-Abad, E. & Moreno, E. NbThermo: a new thermostability database for nanobodies. Database (Oxford) **2023**, baad021 (2023).

100. Strohl, W. R. & Strohl, L. M. Therapeutic antibody engineering: Current and future advances driving the strongest growth area in the pharmaceutical industry. (Woodhead Publishing, 2012).

